# Meta-transcriptomic analysis of virus diversity in urban wild birds with paretic disease

**DOI:** 10.1101/2020.03.07.982207

**Authors:** Wei-Shan Chang, John-Sebastian Eden, Jane Hall, Mang Shi, Karrie Rose, Edward C. Holmes

## Abstract

Wild birds are major natural reservoirs and potential dispersers of a variety of infectious diseases. As such, it is important to determine the diversity of viruses they carry and use this information to help understand the potential risks of spill-over to humans, domestic animals, and other wildlife. We investigated the potential viral causes of paresis in long-standing, but undiagnosed disease syndromes in wild Australian birds. RNA from diseased birds was extracted and pooled based on tissue type, host species and clinical manifestation for metagenomic sequencing. Using a bulk and unbiased meta-transcriptomic approach, combined with careful clinical investigation and histopathology, we identified a number of novel viruses from the families *Astroviridae, Picornaviridae, Polyomaviridae, Paramyxoviridae, Parvoviridae, Flaviviridae,* and *Circoviridae* in common urban wild birds including Australian magpies, magpie lark, pied currawongs, Australian ravens, and rainbow lorikeets. In each case the presence of the virus was confirmed by RT-PCR. These data revealed a number of candidate viral pathogens that may contribute to coronary, skeletal muscle, vascular and neuropathology in birds of the *Corvidae* and *Artamidae* families, and neuropathology in members of the *Psittaculidae*. The existence of such a diverse virome in urban avian species highlights the importance and challenges in elucidating the etiology and ecology of wildlife pathogens in urban environments. This information will be increasingly important for managing disease risks and conducting surveillance for potential viral threats to wildlife, livestock and human health. More broadly, our work shows how meta-transcriptomics brings a new utility to pathogen discovery in wildlife diseases.

**Importance:** Wildlife naturally harbor a diverse array of infectious microorganisms and can be a source of novel diseases in domestic animals and human populations. Using unbiased RNA sequencing we identified highly diverse viruses in native birds in Australian urban environments presenting with paresis. This investigation included the clinical investigation and description of poorly understood recurring syndromes of unknown etiology: clenched claw syndrome, and black and white bird disease. As well as identifying a range of potentially disease-causing viral pathogens, this study describes methods that can effectively and efficiently characterize emergent disease syndromes in free ranging wildlife, and promotes further surveillance for specific potential pathogens of potential conservation and zoonotic concern.

## Introduction

Emerging and re-emerging infectious diseases in humans often originate in wildlife, with free-living birds representing major natural reservoirs and potential dispersers of a variety of zoonotic pathogens (1). Neurological syndromes, such as paresis, are of particular concern as many zoonotic viral pathogens carried by wild birds with the potential to cause neurological disease are also potentially hazardous to poultry, other livestock and humans. Examples of this phenomenon include Newcastle disease virus (NDV, Avulavirus 1; family *Paramyxoviridae*), West Nile virus (WNV, family *Flaviviridae*) and avian influenza viruses (family *Orthomyxoviridae*) (2). Importantly, with growing urban encroachment, the habitats of humans, domestic animals and wildlife increasingly overlap. A major issue for the prevention and control of wildlife and zoonotic diseases is how rapidly and accurately we can identify a pathogen, determine its origin, and institute biosecurity measures to limit cross-species transmission and onward spread. With these ever changing environments, wildlife are also at risk from a conservation perspective, as a number of emerging viral pathogens (WNV, Usutu virus, avian poxvirus, avian influenza, Bellinger River snapping turtle nidovirus) have had adverse population level impacts (3–8). Therefore, a comprehensive understanding of the diversity of the viral community, and the ecology of microbes associated with urban wildlife mass mortality and emergent disease syndromes, will improve our capacity to detect pathogens of concern and lead to improved conservation and public health intervention (3).

Meta-transcriptomic approaches (i.e. total RNA-sequencing) have revolutionized the field of virus discovery, transforming our understanding of the natural virome in vertebrates and invertebrates (9, 10). This method relies on the unbiased sequencing of non-ribosomal RNA, and has been used to identify novel viral species in seemingly healthy animals. In Australia, for example, meta-transcriptomic approaches have been used with invasive cane toads (*Rhinella marina*) (11), waterfowl (12, 13), fish (14) and Tasmanian devils (*Sarcophilus harrisii*) (15). These approaches have also been applied diagnostically in a disease setting, including in domestic animals such as cats (*Felis sylvestris*) (16), dogs (*Canis lupus familiaris*) (17), cattle (*Bos taurus*) with respiratory diseases (18–20), and pythons (*Pythonidae*) with neurological signs (21). Notably, meta-transcriptomics has also been used to identify bacterial diseases, such as tularemia infections in Australian ring-tailed possums (*Pseudocheirus peregrinus*) (22). Hence, metagenomic approaches provide the capacity to rapidly and comprehensively map microbial and viral communities and examine their interactions, improving our understanding of animal health and zoonoses.

Investigations into wildlife diseases are often neglected and under-resourced. Consequently, while many outbreaks and syndromes are reported, until recently limited molecular screening has been performed to characterize the etiology where a novel organism is present. In Australia, several neglected and undiagnosed disease outbreaks have been described in wild avian species including those of suspected viral etiology. Notable examples include two syndromes termed “clenched claw disease” (23–28) and “black and white bird disease” (29–31) that affect rainbow lorikeets (*Trichoglossus moluccanus*) and several species of passerines, respectively.

Clenched claw syndrome (CCS) has been recognized as a form of paresis in rainbow lorikeets in eastern Australia, in which birds present recumbent with poor withdrawal reflexes and clenched feet. Although the syndrome may be multifactorial, a proportion of the cases are characterized by non-suppurative encephalomyelitis and ganglioneuritis, and these cases are suspected to have a viral etiology. Similarly, a series of morbidity and mortality events termed black and white bird disease (BWBD) have occurred in Australian magpies (*Gymnorhina tibicen*), Pied currawongs (*Strepera graculina*), Australian ravens (*Corvus coronoides*) and magpie larks (*Gallina cyanoleuca*) along the Australian east coast (31). Diseased birds present in groups, either dead or paretic. Although these emergent disease syndromes are suspected to be caused by viral infections, they have been poorly described, and to date no viral pathogens have been identified in culture, such that the cause and mechanisms of disease remain elusive.

By exploiting meta-transcriptomic approaches, validated with clinical manifestation and histopathological findings of infection, we investigated the potential viral etiology in historically archived outbreaks of birds fitting the syndrome descriptions associated with black and white bird disease and clenched claw syndrome in Australia, along with other sporadic cases where viral infection was suspected. In particular, we characterized CCS and BWBD and identified a number of novel viruses from the families *Astroviridae, Picornaviridae, Polyomaviridae, Paramyxoviridae, and Parvoviridae* in common urban wild birds.

## Results

### Clinical and histological description of rainbow lorikeets

To formulate a syndrome description for CCS, pathology records from the Australian Registry of Wildlife Health from 451 rainbow lorikeets presenting between 1981 and 2019 were reviewed for unexplained non-suppurative inflammation within the central nervous system. A total of 55 birds were found to match these search terms. The signalments, clinical signs and histological lesions from these birds are summarized chronologically in Table S1. The index case was a juvenile female rainbow lorikeet found in Mosman, NSW in November 1984, and the last known case was recorded in an adult, female from the same location in May 2007. The majority of cases occurred in adult birds (28 adults, 20 juveniles, 7 age unspecified), including 24 males, 21 females and 10 gender unspecified. Although cases were distributed throughout the year, twice as many CCS events occurred in October than any other single month.

Clenched feet was the most prevalent presentation (n = 40), followed by an inability to fly (n = 13), unspecified neurological signs (n = 8), paresis (n = 6), leg paralysis (n = 3), wing paralysis (n = 2), head tilt (n = 3), tremors (n = 3), ascending or progressive neurological signs (n = 2), head bob (n = 2), opisthotonus (n = 1), ataxia (n = 1), and rolling (n = 1) (Figure 1a,b). Body condition was noted in 29 records and 11 birds were classified as being in good condition, 17 birds were considered thin or very thin, and one bird was emaciated. Less common presentations included flying into a window (n = 1) and predation (n = 2). The most common cause of death was euthanasia (n = 37).

**Figure 1.**
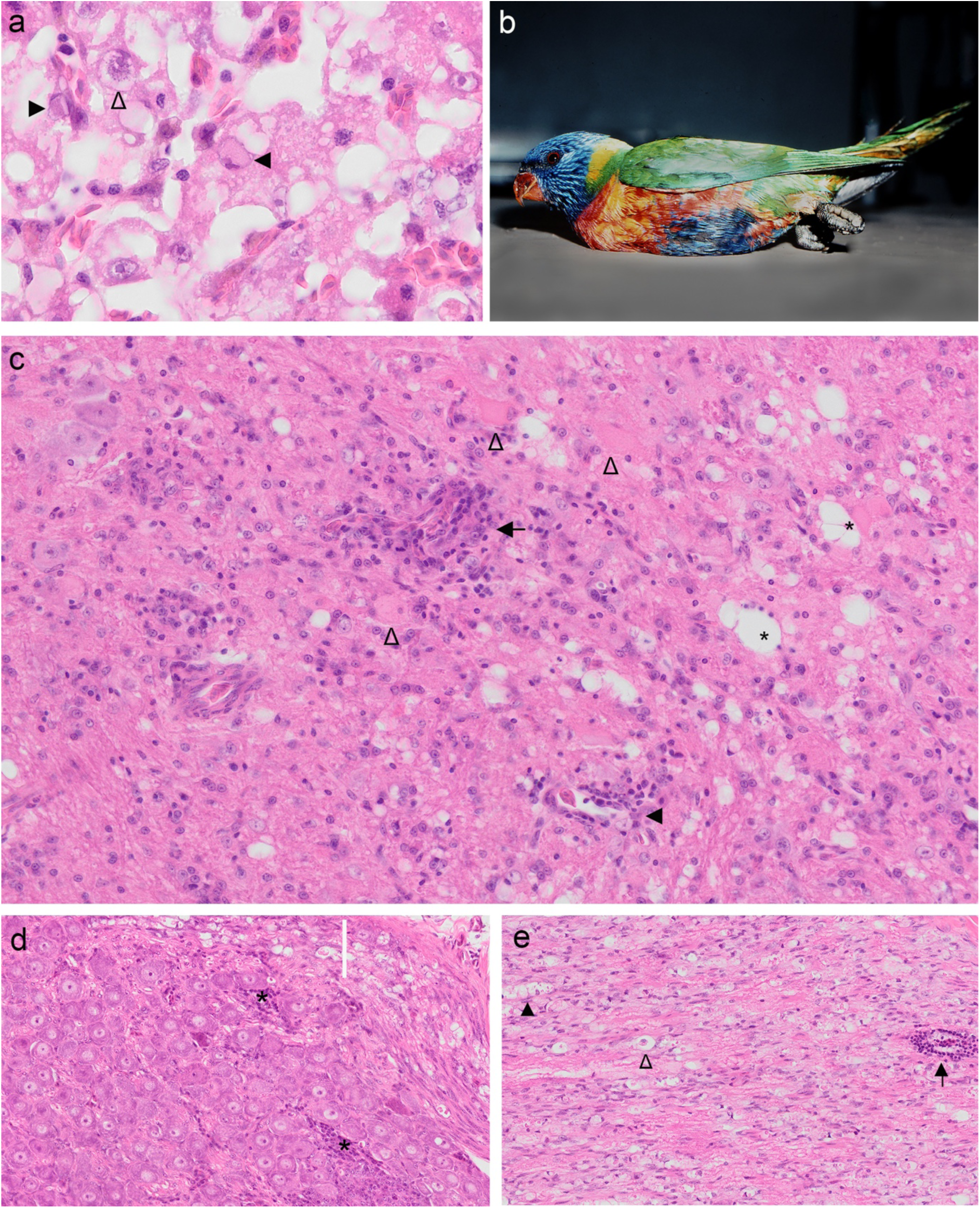
(a) Photomicrograph of rainbow lorikeet hepatocellular intranuclear inclusion bodies (closed arrowheads) and karyomegaly with a stellate chromatin pattern (open arrowhead). (b) Paretic rainbow lorikeet with clenched claws. (c-e) Photomicrographs of lesions characteristic of clench claw syndrome in rainbow lorikeets. Encephalitis within the central cerebellar white matter (c) with perivascular lymphoplasmacytic infiltrates (black arrow), vascular intramural mononuclear cell infiltrates (black arrowhead), gliosis, dilation of axonal chambers (*) and degenerating nerve cell bodies (open arrowheads). Spinal ganglion (d) with multifocal mononuclear cell infiltrates (*), swollen axons and dilated axonal chambers within a white matter tract (white bar). A peripheral nerve (e) with perivascular mononuclear cell infiltration (black arrow), and Wallerian degeneration illustrated by dilated axonal chambers (black arrowhead) and a macrophage within an axonal chamber signifying a digestion chamber (open arrowhead).

Common microscopic lesions in CCS affected lorikeets are illustrated in Figure 1c-e and include mild to severe perivascular cellular infiltrates ranging from one to eight cells deep, composed of lymphocytes, plasma cells and smaller numbers of macrophages. Gliosis, spongiotic change of the neuropil, Wallerian degeneration and nerve cell body degeneration to necrosis were uncommon and restricted to those animals with moderate to severe inflammation. Non-suppurative inflammation within the central nervous system was more prevalent and more severe within the caudal brainstem (28/54), cerebellum (30/54) and spinal cord (44/49), compared with the cerebrum and anterior brain stem (12/54). Peripheral nervous system changes were also noteworthy, as 5 of 47 birds examined had non-suppurative spinal ganglioneuritis, and 18 of 19 birds examined had non-suppurative neuritis, often with Wallerian degeneration (13/19).

An adult, male red-collared lorikeet (*Trichoglossus rubritorquis*) from Queensland was also found with a history of euthanasia following presentation with clenched feet, ataxia, and thin body condition. Histological changes in this animal were consistent with CCS, including non-suppurative lesions that were mild in the cerebrum, and more moderate in the brain stem, cerebellum and spinal cord.

Liver from an adult, male rainbow lorikeet with histological evidence of moderate hepatocellular single cell necrosis, karyomegaly, and unusual amphophilic and variably shaped (stellate, discrete and dense, and ground-glass) intranuclear inclusions (Figure 1a) was included in the meta-transcriptomic investigation based on suspicion of one or more underlying viral infections. This bird presented recumbent, with extensive subcutaneous epithelial lined cysts encapsulating clusters of mites distributed along the wings and cranium, and loss of the distal primary feathers and central tail feathers in a pattern suggestive of psittacine circovirus infection (Registry 4771.1).

### Clinical and histological description of black and white bird disease

A total of 2781 birds in the order Passeriformes were included in the analysis. This identified 51 birds with myocardial degeneration or non-suppurative myocarditis. Two cases were excluded from the analysis, due to incongruity of histological lesions compared with those of all other birds (n = 49) examined. The signalments, clinical signs and histological lesions from these birds are summarized chronologically in Table S2.

Although BWBD was first recognized during an epizootic extending across Victoria, NSW and into Queensland (QLD) in 2006, records from the Australian Registry of Wildlife Health identified the index cases as magpies and currawongs found in Sydney and the NSW central coast in August 2003. The majority of birds examined were magpies (n = 26), currawongs (n = 15), and ravens (n = 6), with adults, juveniles, males and females evenly represented. A solitary adult female figbird and one adult female magpie lark were also among the affected birds. Concurrent mortality was observed in crested pigeons (*Ocyphaps olphotes*), common koels (*Eudynamys scolopacea*) silver gulls (*Larus novaehollandiae*) and indian mynahs (*Acridotheres tristis*). Presentation of affected birds varied, occurring as nine mass morbidity and mortality events, and 25 sporadic cases. The birds examined from outbreaks in 2003, 2006, 2013 and 2015 represent a small fraction of affected birds, and accumulatively total 105 birds reported formally and more than 250 reported informally. Suspected intoxication was a common history in those birds presenting en masse. Although seven birds were found dead or died prior to veterinary examination, clinical signs among the remaining birds included paresis (Figure 2a), recumbency or profound weakness (27), loss of righting reflex or difficulty righting (12), and less commonly dyspnea (4), watery or bloody diarrhea (4), clenched claws (1) and a moribund state (1). Although most birds were recumbent, they were not paralyzed as they had good cloacal tone, withdrawal reflexes, and could stand and shuffle, flap or walk across the room when stimulated. Most ravens had additional clinical signs indicative of central nervous system dysfunction, including ataxia, head tilt, circling, and unusual head posture (4/6). Other species were alert and could accurately grasp items with their beaks, despite being recumbent. The body condition of affected birds varied with 21 noted to be in good body condition, while 27 were classified as thin, very thin or emaciated.

**Figure 2.**
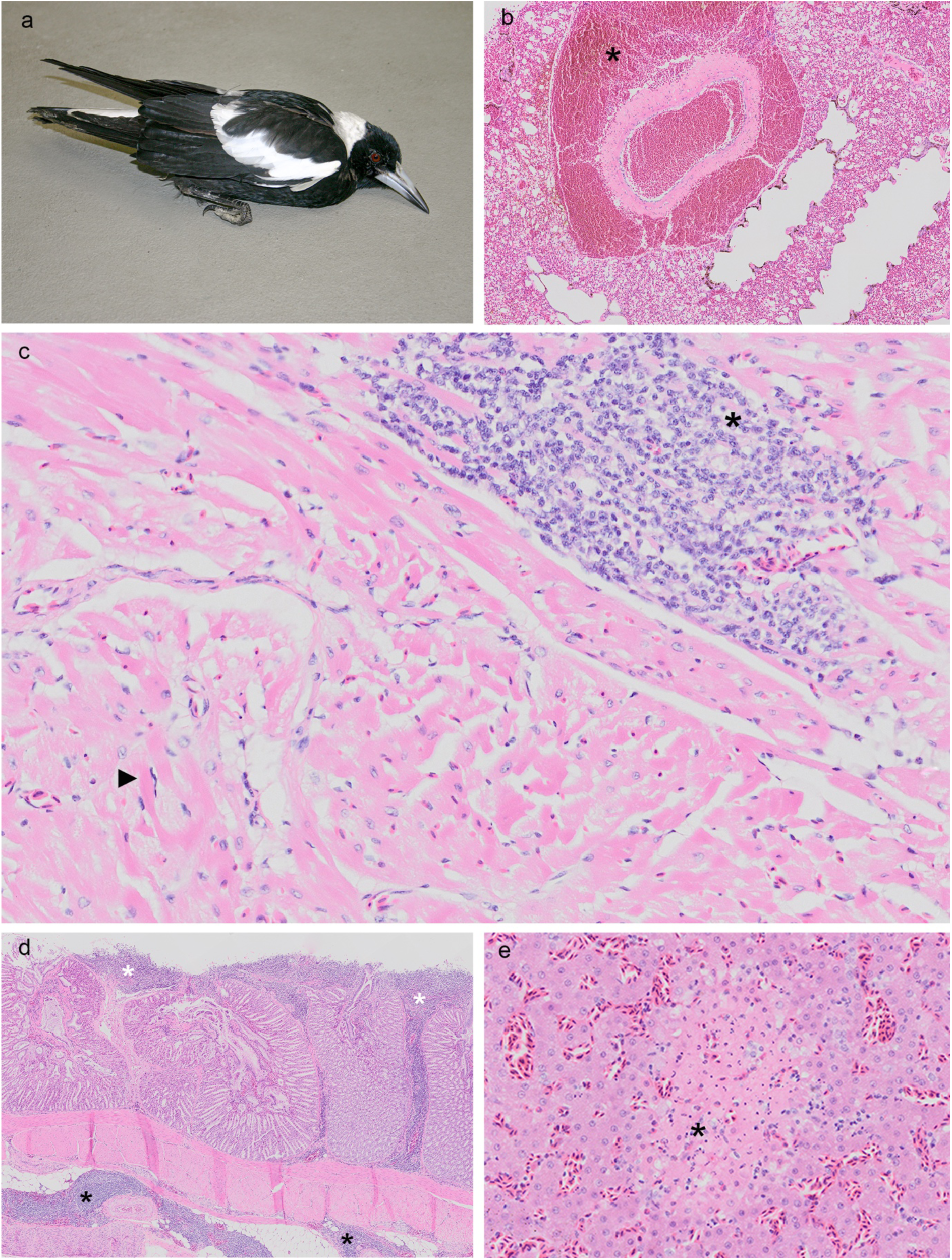
(a) A paretic, but alert, Australian magpie with black and white bird disease. (b-e) Photomicrographs of lesions characteristic of black and white bird disease. Severe perivascular pulmonary hemorrhage (*). Myocardium (c) with a mononuclear cell infiltrate (*) and myocyte degeneration (black arrowhead) signified by hypereosinophilic myofibrils and a pyknotic and peripheralized nucleus. Proventricular mononuclear cell infiltrates (d) within the lamina propria (white *) and surrounding serosal blood vessels (black *). Acute hepatic necrosis (e*).

Gross lesions in affected birds were rare, but included hydropericardium, epicardial hemorrhage, pulmonary congestion to hemorrhage, hemorrhage into the gastrointestinal lumen and fibrinous serositis. Microscopic lesions characteristic of BWBD include the unifying feature of degeneration of cardiac myocytes with or without non-suppurative perivascular and interstitial cardiac inflammation (Figure 2 b-e). Myocyte degeneration or non-suppurative inflammation within skeletal muscle was observed in 24 of the 45 birds examined (four animals were unavailable for examination). Non-suppurative inflammation of the central nervous system was observed in 25 birds. Lesions were most severe in the central cerebellar white matter and included one to four cell deep perivascular cuffs, spongiotic change of the neuropil, degeneration to necrosis of nerve cell bodies and multifocal gliosis. Other common histological features of the syndrome included acute hepatic necrosis or inflammation (25/49), necrosis or non-suppurative gastro-intestinal inflammation (21/49), non-suppurative interstitial nephritis (8/49), mild vasculopathy to marked fibrino-necrotising vasculitis (19/49), non-suppurative perivascular infiltrates throughout serosal surfaces (20/49) and non-suppurative interstitial pancreatitis (11/49). Less common lesions included perivascular pulmonary, cardiac and neural hemorrhage, meningitis and peripheral neuritis. Ravens generally had more severe and widespread central nervous system lesions, and lacked skeletal muscle, renal and serosal inflammation that was evident in the other species. In birds other than ravens, paresis was most likely associated with lesions in striated muscle and the peripheral nervous system. Although Leucocytozoon and microfilaria are common hemoparasites of the bird species examined, and were found in our metagenomic data (32), these organisms were not relevant within disease-related lesions in our cases. Ancillary diagnostic testing for known bacterial and viral pathogens, *Chlamydophyla* species and intoxication revealed no significant findings.

### Meta-transcriptomic virus identification

Archived samples including brain, liver, heart, and kidneys from representative cases of diseased birds were pooled and used to generate nine RNA-sequencing libraries that resulted in 7,725,034 to 26,555,569 paired reads per pool, for meta-transcriptomic analysis. From these data we discovered eight viruses from the RNA virus families, *Astroviridae*, *Paramyxoviridae*, and *Picornaviridae*, the DNA virus families *Adenoviridae*, *Circoviridae*, *Parvoviridae* and *Polyomaviridae*. Two additional viruses - a hepacivirus and a narnavirus - were also identified but unlikely associated with any clinical syndrome.

### Avian avulavirus in brain samples of lorikeets with clenched claw syndrome

The *Paramyxoviridae* are a large group of enveloped linear negative-sense RNA viruses that range from 15.2 to 15.9 kb in length. We identified paramyxovirus-like contigs in the brain libraries of rainbow lorikeets. A RT-PCR designed to amplify the paramyxovirus L protein (301 nt) identified matching RNA in 4/5 brain tissues, corresponding to the four birds presenting with neurological symptoms. Ten overlapping RT-PCRs were then performed to confirm the genomic sequence (Table S4), revealing a typical 3’-N(455 aa)-P(446 aa)-M(366 aa)-F(619 aa)-HN(528 aa)-L(2263)-5’ organization. Based on the phylogenetic analysis of the L and M protein, the novel paramyxovirus identified here - termed Avian paramyxovirus 5/rainbow lorikeet/Australia/CCS - fell into the clade comprising the genus Avulavirus, exhibiting 86.7% and 89.1% amino acid pairwise identity to its closest relative - APMV-5/budgerigar/Japan/TI/75 (GenBank accession number: LC168759).

To provide a provisional assessment of the virulence of this novel virus, we compared F protein cleavage sites between the lorikeet avulavirus and known pathotypes of NDV (Avulavirus 1) that commonly causes disease in avian species, including the velogenic, lentogenic, mesogenic and asymptomatic vaccine types. Specifically, the precursor F glycoprotein (F0) is cleaved into F1 and F2 subunits for the progeny virus to be infectious and to enable replication. The F protein cleavage position for virulent or mesogenic strains contain a furin recognition site comprising multiple basic amino acids. Notably, the lorikeet avulavirus had an F cleavage ‘RRRKKRF’ motif identical to pathogenic NDV strains, suggesting its potential virulence, although this remains to be confirmed (Figure 3).

**Figure 3.**
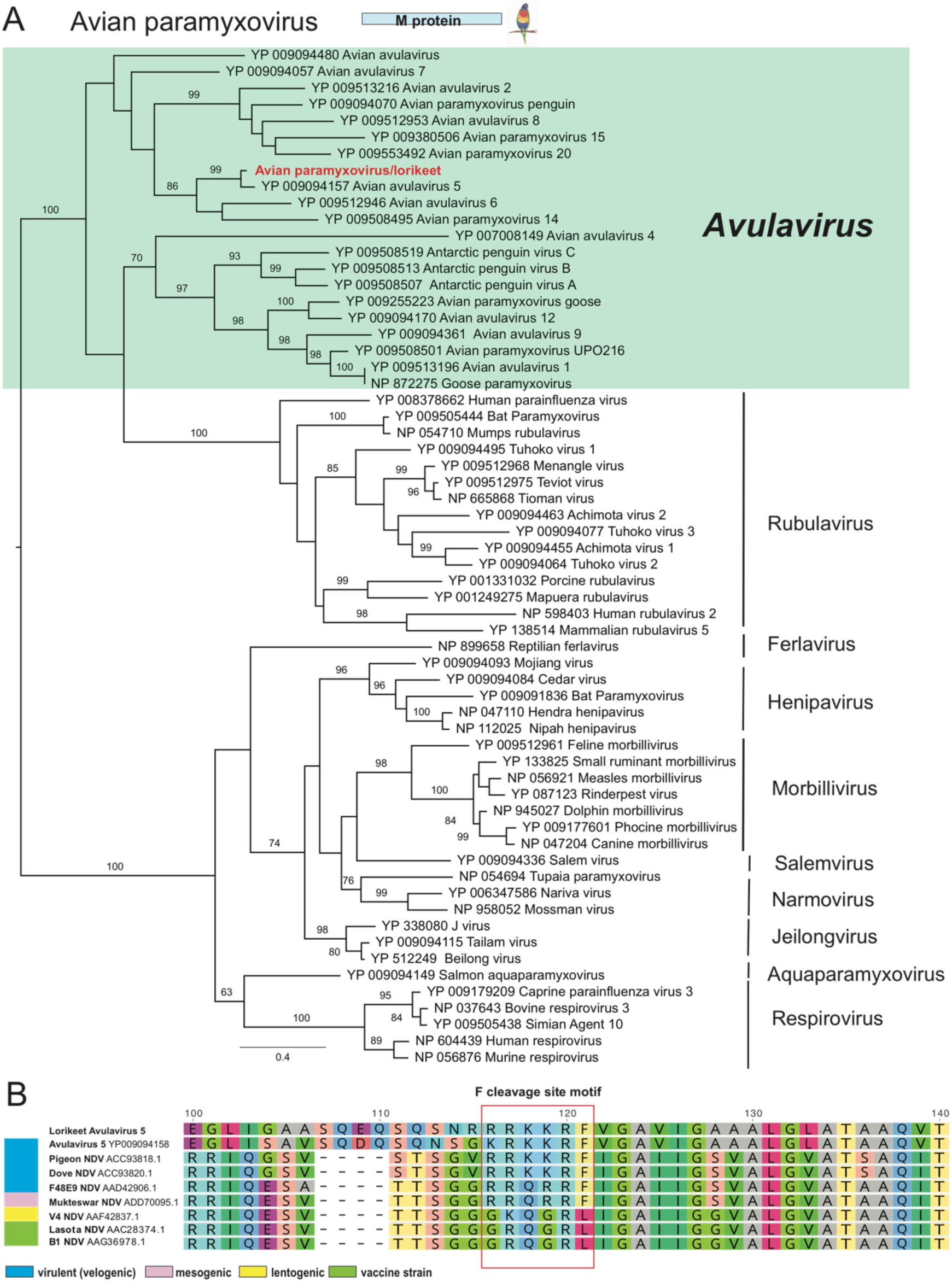
Phylogeny and characterization of a novel avian avulavirus identified from a rainbow lorikeet with clenched claw disease. (a) A maximum likelihood phylogeny of the M gene (366 amino acids) of avian avulavirus plus representative members of the *Paramyxoviridae*, including viruses from the genera *Avulavirus, Rubulavirus, Ferlavirus, Henipavirus, Morbillivirus, Aquaparamyxovirus* and *Respirovirus*. The tree was midpoint rooted for clarity only. The scale bar indicates the number of amino acid substitutions per site. Bootstrap values >70% are shown for key nodes. (b) Characterization of the virulence determination site in the F gene by comparison to representative NDV strains. The typical RRKKR cleavage site motif is highlighted (red rectangle).

### DNA viruses identified in sporadic cases or mortality events rainbow lorikeets

In addition to RNA viruses, we identified three DNA viruses in the context of sporadic mortality events of rainbow lorikeets: avian chapparvovirus, beak and feather disease virus and lorikeet adenovirus. In addition to the RNA-sequencing results, we extracted DNA from corresponding tissues for DNA virus validation through PCR and rolling circle amplification (RCA) assays.

### Lorikeet chapparvovirus

The *Parvoviridae* are a family of small, non-enveloped, dsDNA animal viruses with linear genomes of ∼5 kb in length. We identified parvovirus-like transcripts in both liver RNA-seq libraries from diseased lorikeets. Using RCA to enrich for circular DNAs, we recovered a complete genome of a novel lorikeet chapparvovirus, comprising 4,271 nt with two distinct ORFs that encoded the non-structural protein NS1 (670 aa) and the structural protein VP (542 aa). We further designed specific primers to amplify the targeted VP region (∼248 bp) for screening both the RNA (positive in two liver samples) and DNA (all positive) products. In addition, we inferred two separate phylogenetic trees based on the complete NS1 and VP proteins to determine the evolutionary relationships between the lorikeet virus and other parvoviruses. This revealed that the closest relative to the lorikeet chapparvovirus identified here was an avian-associated red-crown crane parvovirus, yc-9 (GenBank accession number: KY312548.1), although this shares only 48.2% amino acid identity in NS1 and 46.9% identity in VP (Figure 4).

**Figure 4.**
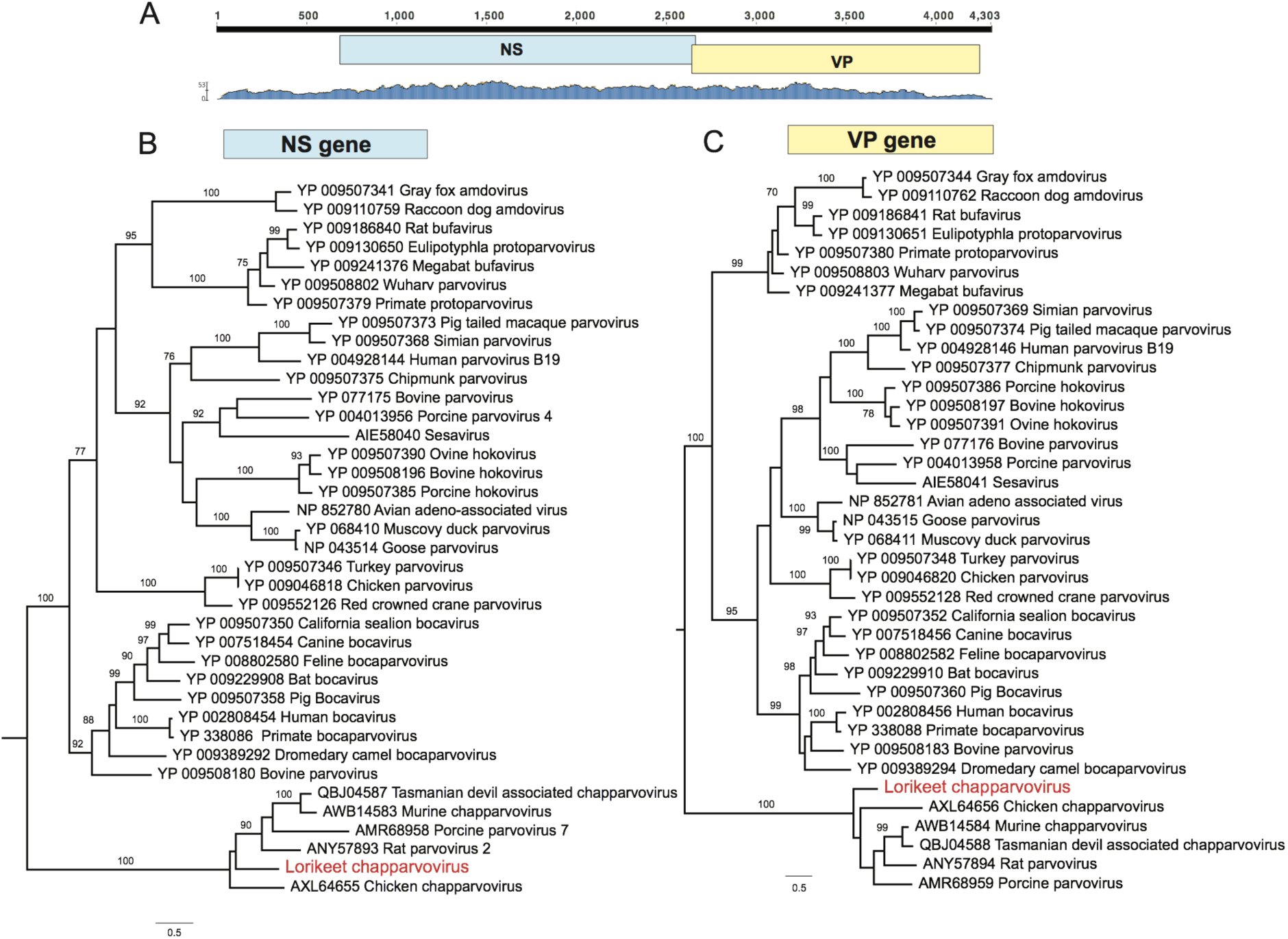
Characterization and phylogeny of the lorikeet chapparvovirus identified here. (b) Schematic representation and read abundance of the genome of lorikeet chapparvovirus. Maximum likelihood phylogenies of the (B) NS gene and (C) VP gene. The tree was midpoint rooted for clarity only. The scale bar indicates the number of amino acid substitutions per site. Bootstrap values >70% are shown for key nodes.

### Beak and feather disease virus

Beak and feather disease virus (BFDV), a member of *Circoviridae*, is highly prevalent in Australian wild birds, particularly psittacine species (33). We identified BFDV-like contigs in all lorikeet RNA libraries, sharing 95% nucleotide identity to known strains of BFDV. Remapping the raw reads to the closest BFDV strain (GenBank accession number: KM887928) gave coverage of the whole virus genome, comprising 2,014 nt, and encoding a replication-associated protein (290 amino acids; aa) and a capsid protein (247 aa) (Figure 5A). We then screened all the RNA samples from rainbow lorikeets using PCR primers targeting the capsid protein (596 nt) (GenBank accession number: KM887928), with the PCR products then Sanger sequenced. The BFDVs identified from individual birds carried distinctive single nucleotide variants, and sequences from each case formed strong phylogenetic clusters suggesting that these results are not due to cross-sample contamination (Figure 5B).

**Figure 5.**
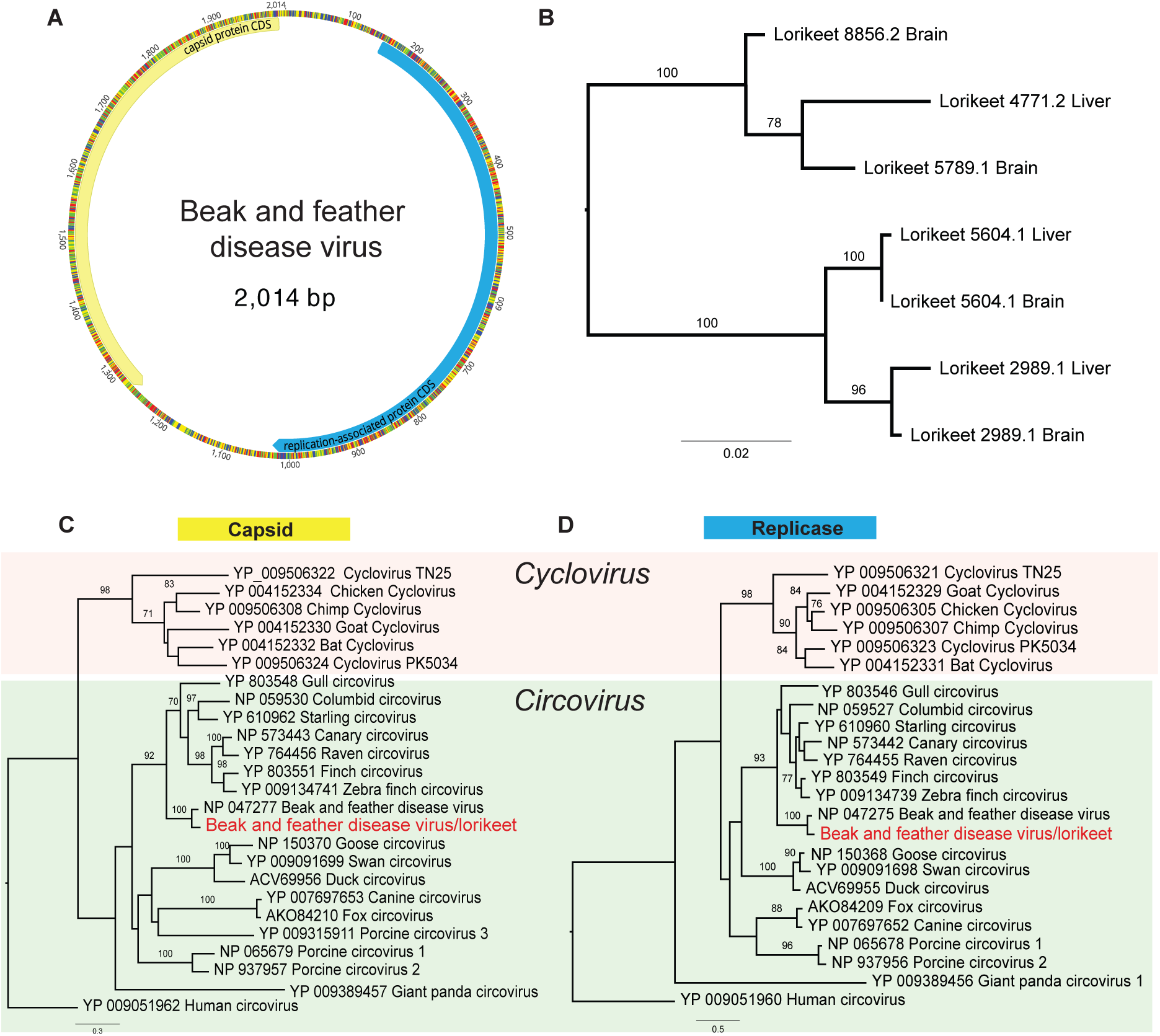
Genomic organization and phylogeny of the beak and feather disease virus (BFDV) identified in this study. (a) BFDV circular genome, annotated with two genes - the capsid protein (Cap) and the replicase-associated protein (replicase). (b) Sanger sequencing of each case detected positive with BFDV. The ML phylogeny of the cap gene (596 nt) was inferred and the BFDVs identified from the same case fell into the corresponding clade. Phylogenetic trees were estimated based on (c) the Cap gene and (d) the Replicase protein. The tree was midpoint rooted for clarity only. The scale bar indicates the number of amino acid substitutions per site. Bootstrap values >70% are shown for key nodes.

### Lorikeet adenovirus

Adenovirus-like contigs were identified from both the brain and liver libraries of rainbow lorikeets. Phylogenetic analyses were performed based on the products of PCR primers designed based on the obtained sequences of polymerase (pol, 843 aa) and hexon (hexon, 580 aa) proteins. The virus identified, termed lorikeet adenovirus (LoAdv), exhibited 76% amino acid identity to the most closely related skua adenovirus (GenBank accession number: YP.0049359313) (Figure 6).

**Figure 6.**
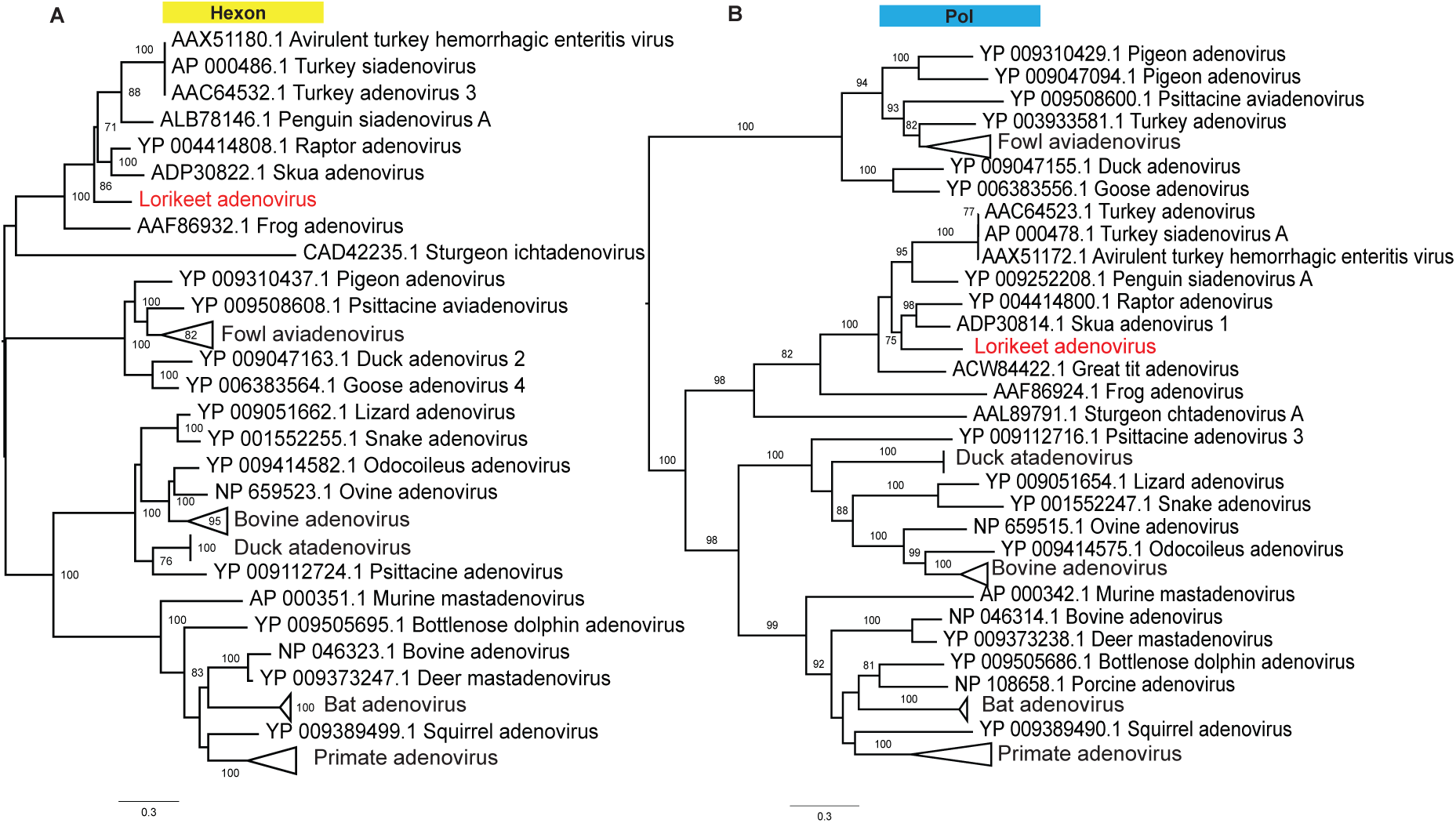
Phylogeny of the novel lorikeet adenovirus identified in this study. ML phylogenies of the (a) Hexon and (b) polymerase protein were inferred with representative members of the *Adenoviridae*. The tree was midpoint rooted for clarity only. The scale bar indicates the number of amino acid substitutions per site. Bootstrap values >70% are shown for key nodes.

### A novel lorikeet hepatovirus in clenched clawed lorikeets

A complete hepatovirus-like (*Picornavirales*) contig was identified in one of the liver libraries of diseased lorikeets, exhibiting relatively high read abundance (∼10,000x coverage depth). The genome of this novel lorikeet hepatovirus, termed lorikeet hepatovirus (LoHepaV), is 7,339 nt in length, and encodes a polyprotein of 2070 amino acids, excluding poly-A tails, that is similar in structure to most Avihepatoviruses (Figure 6). Interestingly, this virus was most closely related to Hepatovirus sp. isolate HepV-bat3206/Hipposideros_armiger/2011 identified from a round-leaf bat, although these two viruses only exhibit 46.3% amino acid identity across the polyprotein. Recently, a lorikeet picornavirus-LoPV-1 (GenBank accession number: MK443503) was identified in fecal samples from rainbow lorikeets in China (34), although this only exhibited 17.4% amino acid identity with the lorikeet hepatovirus identified here (Figure 7).

**Figure 7.**
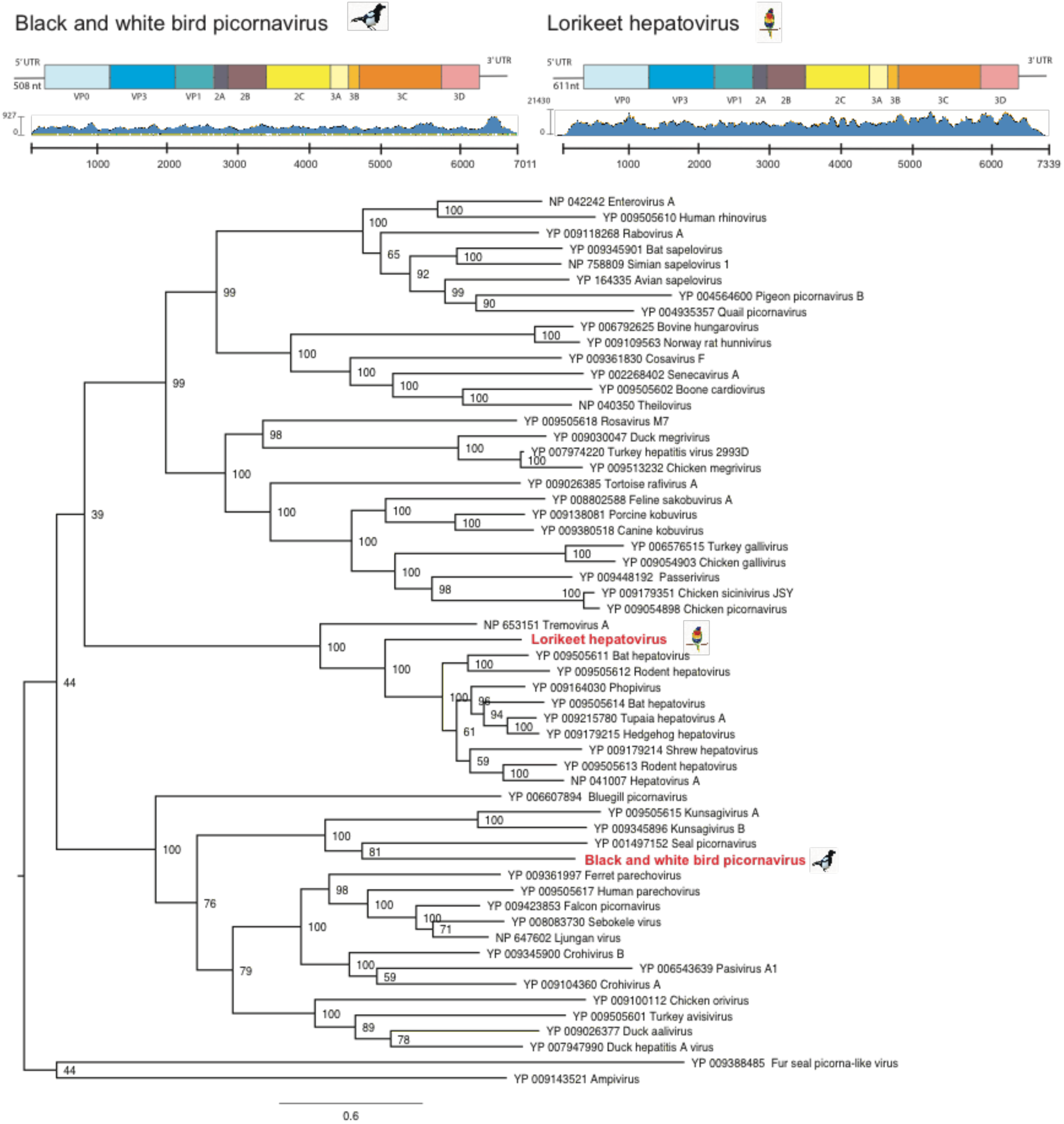
Black and white bird disease picornavirus and lorikeet hepatovirus identified in this study. (a) Schematic representation and read abundance of the black and white picornavirus (upper left) and lorikeet hepatovirus (upper right). (b) ML phylogenetic tree of the polyprotein (containing the RdRp gene) of both viruses. Bootstrap resampling (1,000 replications) was used to assess node support where >70%. GenBank accession numbers are given in the taxa labels. Trees were midpoint rooted for clarify. The scale bars shows the number of amino acid distance in substitution per site.

### Novel picornavirus identified in black and white bird disease cases

The *Picornaviridae* are a large family of single-strand positive-sense RNA viruses, with genomes ranging from 6.7-10.1 kb in length. We identified a highly abundant complete picornavirus-like contig in the library of archived black and white diseases cases. Remapping raw reads to the picornavirus-like transcript showed complete genome coverage (∼300x coverage depth). We named this novel virus black and white bird picornavirus (BWBDpicoV). A RT-PCR targeting this picornaviral contig (∼220 bp) was positive in 2/6 of the brain tissues tested from an Australian magpie (*Cracticus tibicen*) and an Australian raven (*Corvus coronoides*): notably, the two positive birds were those presenting with non-suppurative encephalomyelitis.

BWBDpicoV exhibits classical picornavirus features, with a genome length of 7,011 nt excluding the poly A tails. This genome encodes a polyprotein of 2070 amino acid residues, as well as a 5’ untranslated region (UTR) of 508 nt that includes the putative internal ribosome entry site (IRES). The genome organisation of BWBDpicoV is similar to that of other picornaviruses, with the characteristic gene order: 5′-VP4, VP2, VP3, VP1, 2A, 2B, 2C, 3A, 3B, 3Cpro, 3Dpol-3′ (Figure 7). BWBDpicoV also exhibited such typical picornavirus features as rhv-like domains (111-222,307-424 aa), an RNA-helicase (1172-1271 aa), and an RNA-dependent RNA polymerase (RdRp; 2903-2154 aa). Phylogenetic analysis revealed that BWBDpicoV was most closely related to seal picornavirus (GenBank accession number: NC.009891.1), a member of the genus *Aquamavirus* (Figure 7). However, BWBD picornavirus exhibited low amino acid similarity to the polyproteins of any of these viruses: only 29.5% with Kunsagivirus A (GenBank accession number: YP_009505615.1) and 30.8% with Seal picornavirus type 1 (GenBank accession number: YP_001487152.1). According to the International Committee on Taxonomy of Viruses (ICTV), members of a genus within the *Picornaviridae* should share at least 40% amino acid sequence identity in the polyprotein region. As such, our BWBD may represent a new virus genus.

### A divergent black and white bird astrovirus in a BWBD outbreak from Nowra, NSW

Several astrovirus-like reads were found in the RNA-seq library of the suspected BWBD (i.e. non-suppurative encephalitis of undetermined etiology) outbreak from Nowra, NSW (Table 1). This novel virus, termed black and white bird astrovirus (BWBDastV), was highly divergent in sequence, and contained only 25 reads that covered the partial RdRp of Astrovirus Pygoscelis/DT/2012 (GenBank accession number: KM587711.1) (see below).

**Table 1.**
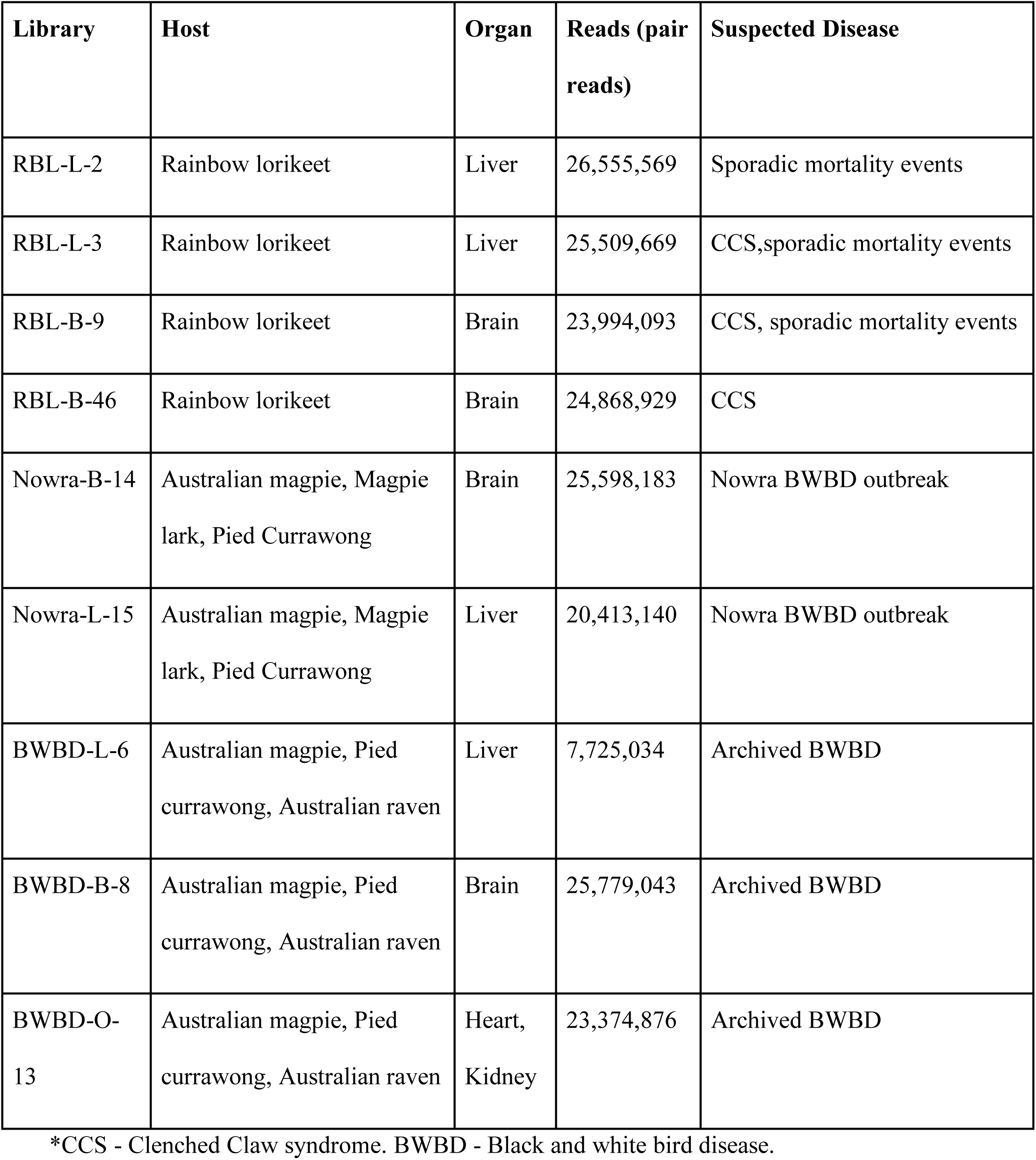
Details of the 9 RNA-seq libraries, the avian species, and associated disease syndrome studied here.

Reads from BWBDastV were used to develop PCR assays to bridge the gap in the RdRp. Using RT-PCR amplification of a 195 nt region targeting the RdRp, we obtained positive hits for BWBDastV in 4 of 5 brain samples. Based on RdRp protein identity and phylogenetic analysis, there was 69.8% amino acid sequence similarity to the closest astrovirus, Astrovirus Pygoscelis/DT/2012 (GenBank accession number: KM587711) sampled from Adelie penguins (*Pygoscelis adeliae*) in Antarctica (Figure 8). Combining the PCR results with clinical and histopathology patterns of diseased birds from the Nowra outbreak, we suggest that BWBDastV might be the cause of the BWBD outbreak (Figure 10B).

**Figure 8.**
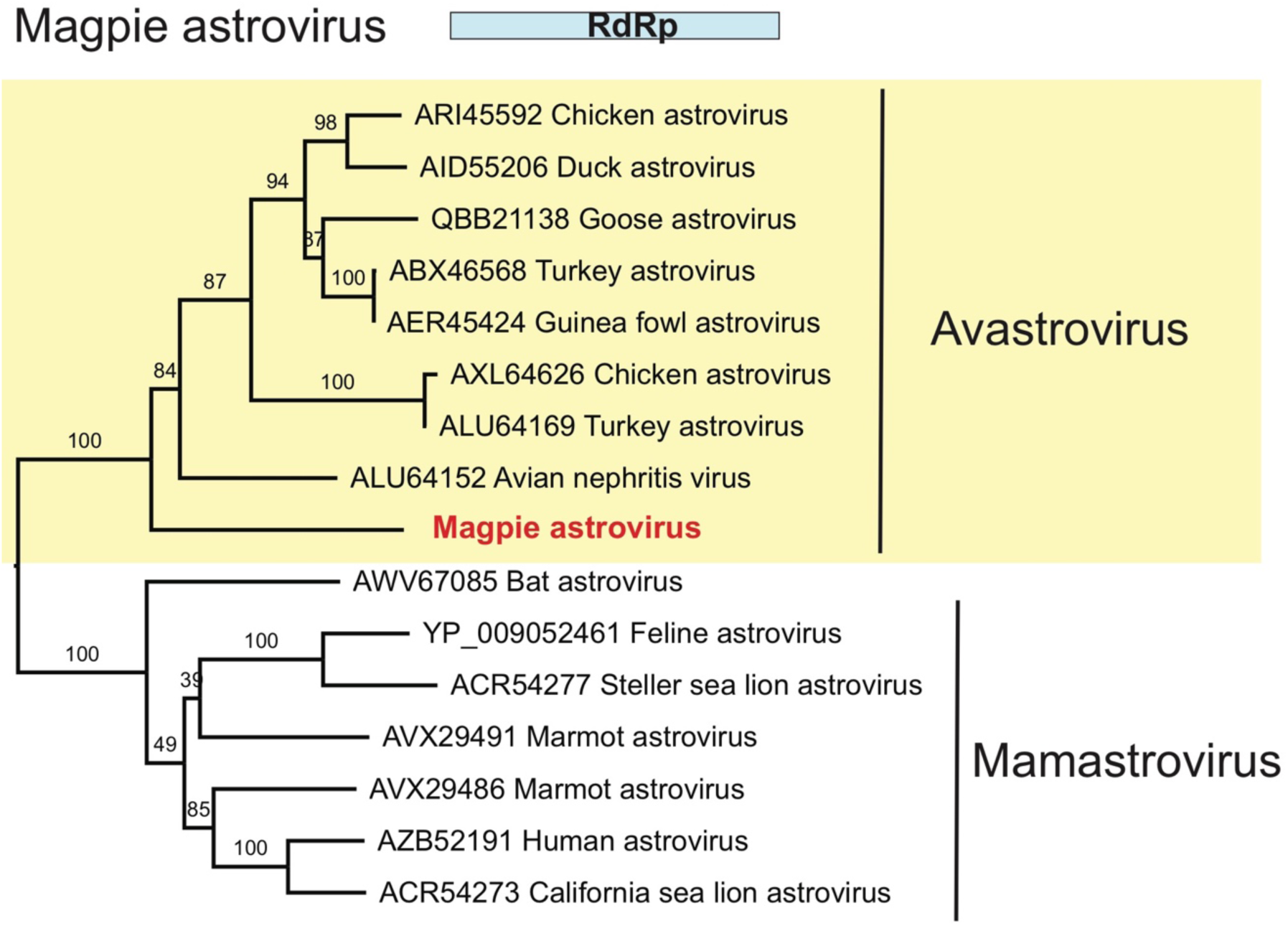
Phylogenetic analysis of magpie astrovirus. ML phylogeny of the partial RdRp gene (267 amino acids) of astroviruses was inferred along with representative members of the *Avastrovirus* and *Mamastrovirus* genera. GenBank accession numbers are given in the taxa labels. Trees were midpoint rooted for clarity only, and branch support was estimated using 1,000 bootstrap replicates, with those >70% shown at the major nodes.

### Novel magpie polyomavirus in the Nowra outbreak

In addition to the novel astrovirus, our RNA-seq analysis identified a novel circular avian polyomavirus in brain and heart tissue from one non-diseased Australian magpie from the Nowra outbreak. Of note, the histopathology of BWBD was not consistent with polyomavirus infection which is not known to be associated with myositis or encephalitis in birds. The genome of the novel magpie polyomavirus - termed magpolyV - comprised 5115 bp with an overall GC content of 46.5%, and we were able to recover full length genome of this virus using RCA and PCR assays. Based on sequence similarity with existing reference polyomavirus proteins we predict that the genome of the novel virus encodes capsid proteins VP1, VP2, VP3 and open reading frame-X (ORF-X) from the late region, as well as two alternative transcripts, large T antigen (LT) and small T antigen (ST) (Figure 9). Consistent with most known polyomaviruses, magpolyV retains the typical conserved motifs and splicing sites in LT, including HPDKGG (DnaJ domain), LRELL and LLGLL (LXXLL-CR1 motif), LFCDE (LXCXE,a pRB1-binding motif) and GAVPEVNLE (ATPase motif). However, the consensus sequence CXCXXC for protein phosphatase 2A binding, mostly found in mammalian polyomaviruses, was not present. Phylogenetic analysis revealed that this magpie polyomavirus consistently clustered within avian lineages, exhibiting 90.3% amino acid its closest relative - butcherbird polyomavirus isolate AWH19840 (accession number: KF360862) - in the LT protein, and 92% amino acid similarity in VP1.

**Figure 9.**
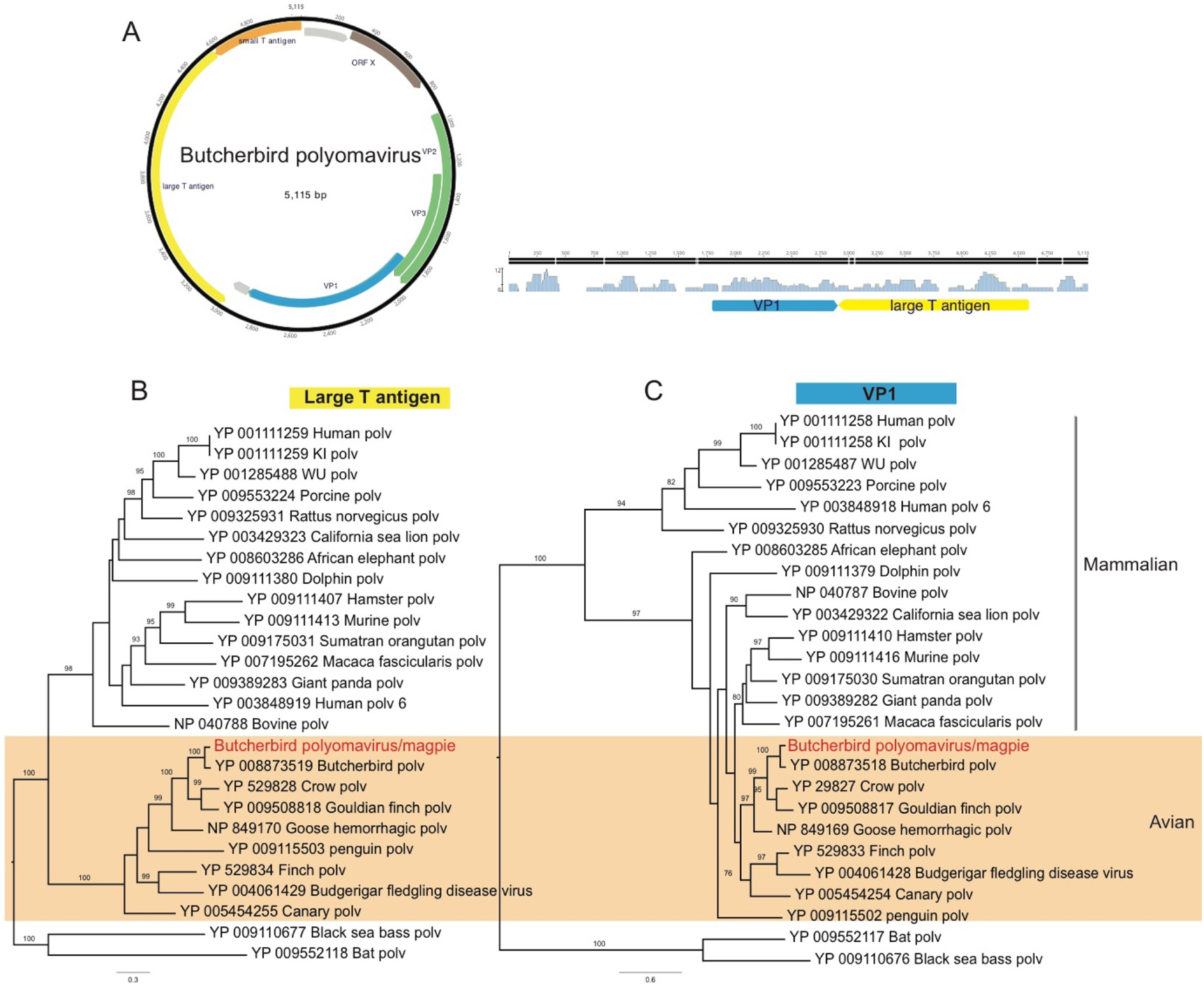
Genome characterization and phylogenetic analysis of the black and white bird polyomavirus identified in this study. (a) Schematic depiction and read abundance of the butcherbird polyomavirus/magpie. ML phylogenetic trees of the (b) large antigen (blue) and (c) VP1 (yellow) protein of the butcherbird polyomavirus/magpie. Bootstrap support (1,000 replicates) was used to assess node support where >70%. GenBank accession numbers are given in the taxa labels. The scale bars show the numbers of substitutions per site.

## Discussion

We characterize the clinical and histological features of CCS and BWBD and apply a meta-transcriptomic approach to identify potential viral etiological agents in wild Australian birds presenting with hepatic inclusion bodies, paresis and non-suppurative myocarditis or encephalomyelitis, elucidating the complex nature of viral disease investigation amid multiple infections. Next-generation metagenomic sequencing, in combination with traditional gross and histological examination, identified several candidate pathogens for disease syndromes that had been reported for decades as undiagnosed, but likely of viral origin. Application of this methodology in the face of emergent disease syndromes or mass mortality events will facilitate the early detection of viruses infecting wildlife, greatly contributing to disease control, conservation of wild populations and the more rapid identification of novel pathogens.

### Clenched claw syndrome

A key element of our paper was the investigation of CCS, a paretic disease of unknown etiology in rainbow lorikeets (*Trichoglossus haematodus*) and a single red-collared lorikeet (*Trichoglossus rubritorquis*) in Australia. The diseased lorikeets present with paresis and were initially often found recumbent or sitting on their hocks with feet clenched, the passive position for the avian foot. Affected birds kept in care progressively developed head tilt, ataxia, and paralysis. The disease syndrome has primarily occurred along coastal New South Wales and South-East Queensland since the early 1980s. Although previous studies report 5-10% of free-living rainbow lorikeets rescued annually presenting with this syndrome, it seems likely that the clinical syndrome in fact encompasses two or more disease etiologies (35). For example, lead poisoning and plant based intoxication have been proposed as contributing to syndromes described as clenched claw or drunken lorikeet syndrome (36). We focused on the investigation of 55 cases of CCS characterized by non-suppurative encephalomyelitis and often ganglioneuritis in which a viral etiology has been suspected. The meta-transcriptomic results, in conjunction with the clinical signs, histopathologic data and confirmation through PCR assays, suggest that the Avulavirus (APMV-5) played a potential leading role in the pathogenesis of CCS in these diseased birds (Figure 10a). An increasing incidence of clinical AMPV infection in wild birds is reported worldwide (37). APMVs have been categorized into 12 serotypes, and APMV1 (NDV) causes significant economic impact on the poultry industry as well as population level impacts on wild birds (37). Both NDV and APMV5 have been reportedly associated with disease outbreaks where mortality rates approach 100%, although the pathogenicity of APMV5 varies substantially among aviary species. APMV5 was first isolated from a fatal outbreak of caged budgerigars in Japan in 1974, followed by epizootic outbreaks in budgerigars in the UK (38) and Australia (39). Interestingly, APMV5 will not replicate in chicken embryonated eggs, it lacks a virion hemagglutinin, and it is not pathogenic to chickens (40). Phylogenetically, the AMPV5 identified here is closely related to the Brisbane and Japanese strains identified from budgerigars (41). Because of the possibility of cross-species transmission, the prevalence and pathogenicity of avian paramyxoviruses circulating among wild birds in Australia clearly merits further investigation.

**Figure 10.**
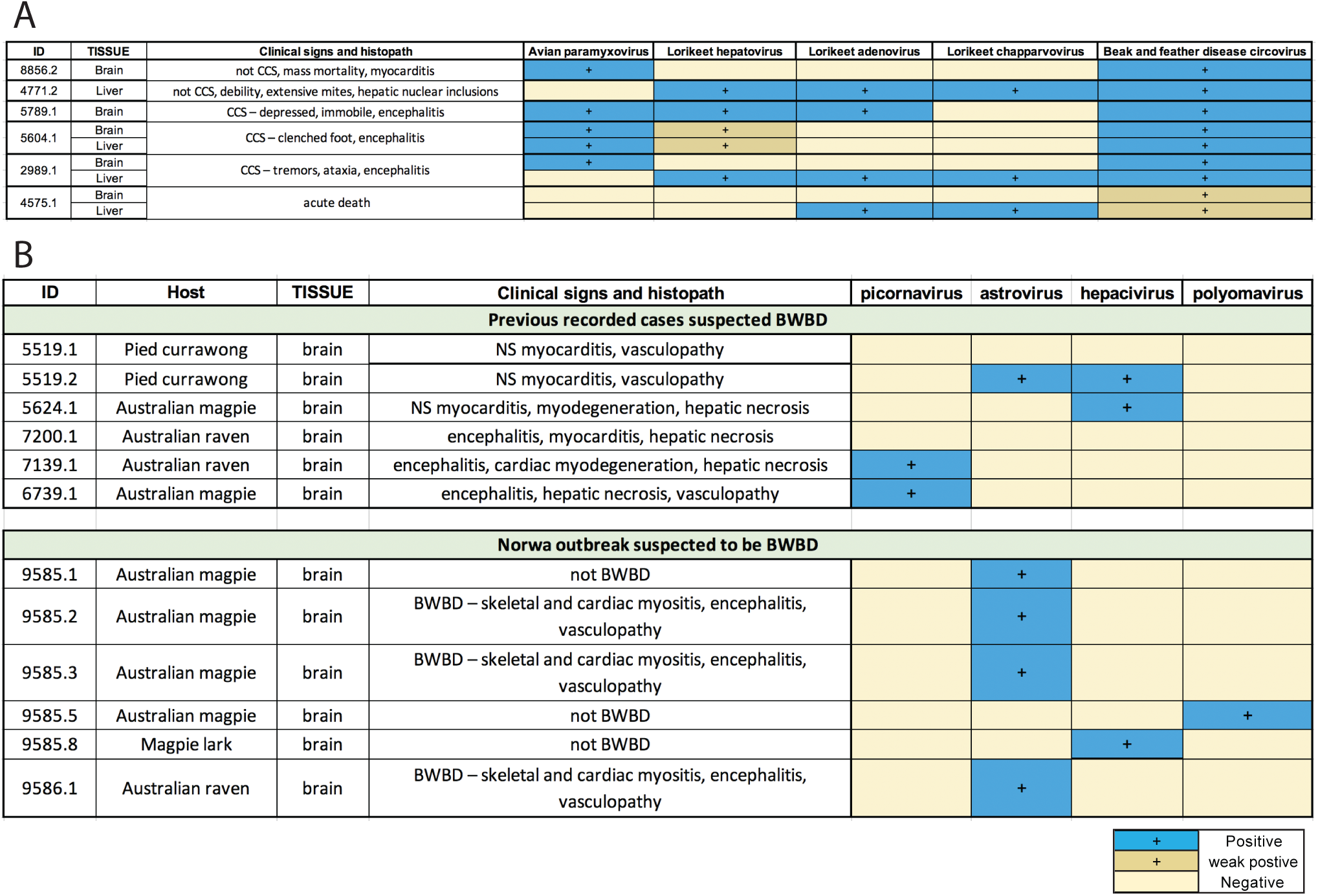
Summary of cases with meta-transcriptomic pathogen identification and confirmation through PCR assays. (a) Clenched claw syndrome. (b) Black and white bird disease. Blue: PCR positive, yellow: negative, dark yellow: weak positive, NS: non-suppurative, CCS: Clenched claw syndrome, BWBD: Black and white bird disease.

### Detection of avian chapparvovirus, avian adenovirus and beak and feather disease viruses in rainbow lorikeets

Chapparvoviruses are being increasingly identified in metagenomic studies, and seemingly have a wide host range (42), including bats (43), and the feces of turkeys (44), red-crowned cranes (45), wild rats (46) and pigs (47). Importantly, a murine kidney parvovirus was recently found in immunodeficient laboratory mice strains, causing renal failure and kidney fibrosis (48), indicating that some chapparvoviruses are associated with highly virulent infections.

Avian polyomavirus, beak and feather disease virus, and avian adenovirus appear to be prevalent in psittacine species in Australia (49), compatible with the data presented here. As multi-viral infection is common, our analysis of viral communities associated with neglected avian diseases enabled a comprehensive understanding beyond the one-host, one-virus system (13). Many known viral infections are relevant clinical issues in psittacine species due to their association with acute death, disease, the capacity to induce immunocompromise, and difficulties in treatment and control (50). These have been associated with a variety of RNA viruses including avian paramyxovirus (37), avian influenza virus (51, 52), and avian bornavirus (53), and DNA viruses such as psittacine beak and feather disease virus (PBFD) (54), avian polyomavirus infection (APV) (55), psittacid herpesvirus (PsHV) (56), psittacine adenovirus (PsAdV) (57), poxvirus infection (58), and papillomavirus infection (59, 60).

### The role of astroviruses, polyomaviruses and picornaviruses in black and white bird disease

Viruses of the family *Astroviridae* comprise genomes of 6.8-7kb, infect multiple mammalian species including humans, and also avian species. Astrovirus infection is most commonly associated with gastroenteritis, and rarely neurological symptoms. A number of novel divergent novel astroviruses in wild birds have recently been discovered through metagenomic surveillance (61–63). Many avian astroviruses, including avian nephritis virus, duck and turkey astroviruses and duck hepatitis virus 2, have considerable economic impact on the poultry industry and these viruses are widely distributed in wild birds (62). Despite this, the diversity and ecology of these viruses, including their potential interspecies transmission events, are largely unknown. Previously, astrovirus-associated central nervous system impairment had been reported in mink, human, bovine, ovine, and swine (64–68).

In mammals such as dogs and cats, astroviruses are commonly isolated with enteric bacterial pathogens in sporadic gastroenteritis outbreaks (69), while in birds, astrovirus infection manifests as a broader disease spectrum including enteritis, hepatitis and nephritis, but rarely neurological symptoms. The astrovirus identified here (BWBDastV) was present in each bird with non-suppurative encephalomyelitis, concurrent with myositis, coelomitis and myocarditis from a single outbreak. However, formal classification of the virus will require additional sequence information. In addition, further surveillance and *in situ* diagnostic modalities are required to reveal the association between BWBDastV, clinical illness of the birds, and the histological lesions observed.

A novel polyomavirus - magpolyV - was identified from the brain and heart of one of the non-diseased Australian magpies (*Cracticus tibicen*) that died in the same BWBD outbreak. An avian polyomavirus (APV) - budgerigar fledgling disease polyomavirus – has been documented in young budgerigars and other psittacine species, causing feather abnormalities, abdominal distension, head tremors and skin hemorrhages, reaching infection rates of up to 100% in aviaries worldwide (70). The clinical presentation and degree of susceptibility to APV infection varies from skin diseases to acute death. In the case of non-psittacine species, goose polyomavirus has been characterized as the etiologic agent of fatal hemorrhagic nephritis and enteritis of European geese (HNEG) (71). Other species of polyomaviruses have been identified in finches (72), crows (73) butcherbirds (74), and a novel APV was recently associated with a fatal outbreak in canaries (75). Moreover, a recent study revealed an APV isolated from pigeon feces in China showing almost 99% identity with previously identified psittacine strains, suggesting a much broader host range of APVs and undermined cross-species transmission (76).

Picornaviruses are commonly identified infecting a variety of avian species in metagenomic studies. In the context of overt disease, notable avian picornaviruses include avian encephalomyelitis viruses in chickens, pheasants, turkeys (77), duck hepatitis A in ducklings (78), and a novel poecivirus that has been identified strongly associated with Avian keratin disorder (AKD) in Alaskan birds (79, 80). We identified two novel picornaviruses in this study - black and white bird picornavirus and a lorikeet hepatovirus - the pathogenic potential of which merits further investigation. Interestingly, a novel lorikeet picornavirus (LoPv-1, GenBank accession number MK443503) was identified in the feces of healthy rainbow lorikeets (*Trichoglossus haematodus*) in China (34), although the polyprotein only shared 12.6% similarity with the lorikeet hepatovirus identified here.

The high frequency of microbial co-infections suggests that rather than being the primary cause of the disease outbreak, some novel avian viruses might augment another pathogen, causing immunosuppression in immuno-compromised hosts (81). Although we were successful in elucidating seven candidate viral pathogens that eluded detection through viral culture, it is clear that additional experimental evidence such as *in vivo* inoculation experiments and *in situ* methodologies to identify pathogens within lesions are critical for clarifying the relationship between infection and the emergent disease syndromes we consider. Notably, the metagenomic approach utilized here is not limited to viruses and can be useful in the detection of bacteria, fungi, protozoan and metazoan parasites. For example, the same data set provided a meta-transcriptomic signal of Leucocytozoon manifestation in BWBD affected birds (32), although our histological data suggests that this was not a primary or secondary pathogen.

The utility of meta-transcriptomics in the context of a thorough and multi-disciplinary diagnostic approach is to rapidly identify viruses and other microbial species, including those that are highly divergent, with greater sensitivity and capability than conventional diagnostic tests. The sequencing data and PCR assays developed here enhance diagnostic capabilities for avian disease and enable epidemiological surveillance to better understand the ecology and impacts of the viruses described. More broadly, our study highlights the value of proactive viral discovery in wildlife, and targeted surveillance in response to an emerging infectious disease event that might be associated with veterinary and public health.

## Materials and methods

### Animal ethics

Wild birds were examined under the auspices of NSW Office of Environment and Heritage Licenses to Rehabilitate Injured, Sick or Orphaned Protected Wildlife (#MWL000100542). Diagnostic specimens from recently deceased birds were collected under the approval of Taronga Animal Ethics Committee’s Opportunistic Sample Collection Program, pursuant to NSW Office of Environment and Heritage issued scientific licenses #SL10469 and SL100104.

### Sample collection

Samples were collected between 2002 and 2013 from 18 birds, predominantly from within the Sydney basin and the coastal center of NSW. Fresh portions of brain, liver, heart, and kidney were collected aseptically and frozen at −80°C. Additionally, a range of tissues were fixed in 10% neutral buffered formalin, processed in ethanol, embedded with paraffin, sectioned, stained with hematoxylin and eosin, and mounted with a cover slip prior to examination by light microscopy. Giemsa stains were applied to a subset of tissue samples to determine whether protozoa were present within lesions.

### Historical Case Review

The records of the Australian Registry of Wildlife Health dating back to 1981 were interrogated to identify rainbow lorikeets with clinical signs relating to the nervous system, and those with unexplained non-suppurative inflammation in the central nervous system. An additional search was conducted to identify passerine birds with unexplained non-suppurative inflammation within cardiac or skeletal muscle. Retrieved records were investigated to determine the signalment, clinical signs and histological lesions of affected animals.

The severity of non-suppurative inflammation was graded on a scale of 0-4, where 0 indicated no discernible lesions and 4 represented severe and extensive mononuclear cell inflammation. Degeneration and necrosis were graded on a scale of 0-4 where 1 represented mild, multifocal single cell degeneration or necrosis, and 4 characterized extensive coagulative necrosis or malacia. Wallerian degeneration of the white matter tracts within the brain, spinal cord, central and peripheral nerves was noted when present. In rainbow lorikeets the lesions were graded in the cerebrum, brain stem, cerebellum, spinal cord, ganglia, and peripheral nerves. Whereas in the passerines, degeneration or necrosis, and non-suppurative inflammation were graded in skeletal and cardiac muscle, brain, liver, gastrointestinal tract, pancreas, blood vessel walls, and renal interstitium.

Avian influenza and NDV were excluded using real-time PCR on oropharyngeal, conjunctival and cloacal swabs from 22 magpies and 12 other wild birds (including currawongs, magpies and ravens) (Flu DETECT, Zoetis). Serological testing of plasma from the same birds comprised hemagglutination inhibition to identify NDV antibodies, and competitive ELISA tests targeting Avian influenza virus and flavivirus antibodies. Sixty-eight tissue samples from a sub-group of 13 magpies, currawongs and ravens identified as having acute histological lesions suggestive of viral infection were subjected to further avian influenza, NDV and WNV specific rt-PCR and virus isolation. Virus isolation was attempted in both mammalian and insect cell lines (Peter Kirland pers comm.). Additional viral culture was attempted on the same samples inoculated into chick embryos while assessing hemagglutinating agents, and in Vero cells assessing cytopathic effect consistent with WNV infection. Clearview antigen capture ELISA (Unipath, Mountain View, Calif.) targeting *Chlamydophila* species was conducted on splenic tissue from eight birds with BWBD (magpies, currawongs and a raven and magpie lark). Liver and lung tissues from eight magpies and currawongs were subjected to routine aerobic and anaerobic bacterial and fungal culture. Finally, liver tissue from eight magpies and currawongs, representing the 2006 and 2015 BWBD epizootics, were subjected to toxicological testing to exclude the presence of a variety of acaricides, fungicides, organophosphates, carbamates, synthetic pyrethroids, and organochlorines and their metabolites.

### RNA extraction, library construction and sequencing

Viral RNA was extracted from the brain and liver, heart and kidney samples of animals using the RNeasy Plus Mini Kit (Qiagen, Germany). RNA concentration and integrity were determined using a NanoDrop spectrophotometer (Thermo) and TapeStation (Agilent). RNA samples were then pooled in equal proportions based on animal tissue type and syndrome. Illumina RNA libraries were prepared on the pooled samples following rRNA depletion using a RiboZero Gold kit (Epidemiology) at the Australian Genome Research Facility (AGRF), Melbourne. The rRNA depleted libraries were then sequenced on an Illumina HiSeq 2500 system (paired 100 nt reads)

### Virome meta-transcriptomics

Unbiased sequencing of RNA aliquots extracted from diseased animals in each outbreak were pooled based on host species, clinical syndrome and histological findings as shown in Table 1. The RNA sequencing reads were trimmed of low quality bases and any adapter sequences before *de novo* assembly using Trinity 2.1.1 (82). The assembled sequence contigs were annotated using both nucleotide and protein BLAST searches against the NCBI non-redundant sequence database. To identify low abundance organisms, the sequence reads were also annotated directly using a BlastX search against the NCBI virus RefSeq viral protein database using Diamond (83) with an e-value cutoff of <10^−5^. Open reading frames were then predicted from the viral contigs in Geneious v11.1.2 (84) with gene annotation and functional predictions made against the Conserved domain databases (CDD) (85). All sequences were aligned using the E-INS-i algorithm in MAFFT version 7 (86). Virus abundance was assessed using a read mapping approach, in which reads were mapped to the assembled viral contigs using the BBmap program (87).

### Rolling-circle amplification assays for circular DNA viruses

To recover the complete genomes of the polyomaviruses and circoviruses identified through our RNA-Seq analysis, we enriched for circular DNA using rolling-circle amplification (RCA) (88). Briefly, genomic DNA extracted from animal tissue and combined with a reaction buffer containing random primers, 1X phi29 Buffer, DTT, Bovine Serum Albumin, dNTPs, and phi29 DNA polymerase (Thermo Scientific, Australia) and then incubated at 30°C for 16h. The RCA products were then purified using the Monarch PCR & DNA Cleanup Kit (NEB), then quantified using the Qubit dsDNA broad range assay. Nextera XT DNA libraries were then prepared and sequenced on an Illumina MiSeq to a depth of ∼2M reads (2 × 150nt).

### RT-PCR assays and Sanger sequencing

Total liver and brain RNA from individual birds was reverse transcribed with SuperScript IV VILO Mastermix (Invitrogen). The cDNA generated from the sampled tissues was used for viral specific PCRs targeting regions identified by RNA-Seq. RT-PCR primers were designed to bridge any genomic gaps based on detected transcripts of the paramyxoviruses, polyomaviruses, adenoviruses, and astroviruses identified in the RNA-seq data (Tables S3 and S4). Accordingly, all PCRs were performed using Platinum SuperFi DNA polymerase (Invitrogen) with a final concentration of 0.2 µM for both forward and reverse primers. All PCR products were visualized by agarose gel electrophoresis and Sanger sequencing. Long RT-PCR products were also sequenced using Nextera XT and MiSeq, as per RCA assays above.

### Phylogenetic analysis

Conserved domains within the viral proteins produced were used in phylogenetic analyses to determine the evolutionary relationships of the viruses identified here. Specifically, in the case of RNA viruses we used amino acid sequences of the RNA-dependent RNA polymerase (RdRp) that is the most conserved protein among this group. After removing all ambiguously aligned regions using TrimAl (89), phylogenetic trees were inferred using the maximum likelihood method (ML) implemented in PhyML version 3.0 (90), employing a Subtree Pruning and Regrafting topology searching algorithm and the LG model of amino acid substitution. Bootstrap resampling with 1000 replications under the same substitution model was used to assess nodal support. All phylogenetic trees were then visualized using FigTree v1.4.3 (http://tree.bio.ed.ac.uk/software/figtree).

### Nucleotide Sequence Accession Numbers

The RNA sequencing data in this study has been deposited in the GenBank Sequence Reads Archive under accession numbers PRJNAXXXX. All genome sequences of identified viruses have been uploaded in GenBank under accession numbers XXXXX to XXXXXX.

## Acknowledgements

This research was supported by the Taronga Conservation Society Australia, Taronga Conservation Science Initiative, and New South Wales National Parks and Wildlife Service. ECH is supported by an ARC Australian Laureate Fellowship (FL170100022). We thank Drs. Bill Hartley, Rod Reece, Cheryl Sangster, Shannon Donahoe, Richard Montali, and Cathy Shilton for their diagnostic contributions on individual cases. Virology personnel at the NSW Department of Planning, Industry and Environment’s Elizabeth Macarthur Agricultural Institute, and CSIRO’s Australian Animal Health Laboratories are acknowledged for their considerable body of work pursuing viral culture in the BWBD investigation.

## Supplementary Information

**Table S1.** Presentation and pathology of rainbow lorikeets with clench-claw syndrome.

**Table S2.** Presentation and pathology of passerines with myocardial degeneration and myocarditis.

**Table S3.** PCR primers used to amplify viral sequences from bird tissues.

**Table S4.** Primers used to recover full length genome of the viruses identified.

