## Supplementary Information for "Meta-transcriptomic analysis of virus diversity in urban wild birds with paretic disease"

**Supplementary Table 1.** Presentation and pathology of Rainbow Lorikeets with Clenched Claw Syndrome

| | | | | Degree of Non-suppurative Inflammation | | | | | |
|---|---|---|---|---|---|---|---|---|---|
| **Case ID** | **Date** | **Signalment** | **Presentation** | **Cerebrum** | **Brain stem** | **Cerebellum** | **Spinal cord** | **Ganglia** | **Nerve** |
| U88 | 19-11-84 | J/F | Clenched feet, paresis, ascending neurological signs. Euthanised. | 1 | 2/N1 | 1 | 1 | - | - |
| U131 | 28-10-85 | J/- | clenched foot (1), unable to fly, wing tremors, weakness. Died. | 0 | 1/N1 | 0 | 2/N2 | 0 | - |
| U147 | 28-12-85 | J/M | Clenched feet, leg and wing paralysis. Very thin. Moribund. | 1 | 2/N1 | 2/N1 | 2/N2 | 0 | 0 |
| U150 | 10-01-86 | J/F | Clenched feet. Leg paresis. Wings and head normal. | 2 | 2 | 2 | 2/N2 | 0 | - |
| U205 | 11-03-86 | A/F | Clenched feet, unable to fly, prostrate. | 0 | 0 | 0 | 2/N2 | 2 | 2/WD |
| U270 | 13-05-86 | A/M | Leg paralysis. Died. | 0 | 0 | 0 | 2/N1 | 0 | WD |
| U309 | 25-08-86 | A/F | Clenched feet, unable to fly, alert. Euthanasia. | 0 | 0 | 0 | 1 | 0 | - |
| U339 | 20-10-86 | J/M | Clenched feet. | 0 | 2 | 2 | 0 | 0 | - |
| U340 | 20-10-86 | J/F | Clenched feet. | 1 | 1 | 1 | 1 | 0 | - |
| U342 | 21-10-86 | A/M | Clenched feet, eating well, unable to fly or perch. | 2 | 2 | 2 | 2/N2 | 0 | WD |
| U343 | 21-10-86 | A/- | Clenched feet, unable to fly. | 0 | 0 | 0 | 2/N1 | | |
| U345 | 25-10-86 | A/M | Clenched feet, head tilt, unable to fly or walk. Euthanised. Good condition. | 0 | 2 | 0 | 0 | 0 | - |
| U379 | 01-12-86 | A/M | Clenched feet, unable to fly. Euthanised. Good condition. | 0 | 0/N1 | 0 | 0/N1 | 0 | WD |
| U517 | 19-06-87 | A/M | Clenched feet, progressive neurological signs. Euthanised. Good condition. | 0 | 0 | 2/N1 | 2/N1 | 0 | - |
| U559 | 10-09-87 | J/F | Clenched feet, ataxic, head down. Euthanised. Thin. | 1 | 1 | 2 | 1 | 0 | - |
| U586 | 05-10-87 | J/F | Flew into window. Neurological signs. Euthanised. | 0 | 3 | 3 | 2 | 0 | - |
| U584 | 28-09-87 | A/M | Head bobbing, weak, weak neck. Perching. | 0 | 3 | 3 | 3 | 0 | - |
| U594 | 21-10-87 | J/M | Head nodding. Rolling over. Neurological signs. Euthanasia. | 0 | 2 | 2 | 1 | 0 | - |
| U598 | 19-10-87 | J/M | Neurological signs. Euthanised. | 0 | 2 | 2 | 0/WD1 | 0 | - |
| U645 | 29-12-97 | J/M | Unable to fly then neurological signs. Euthanised. | 0 | 0 | 2/N1 | 0 | 0 | - |
| U654 | 22-01-88 | A/F | Clenched feet, leg paresis. Euthanised Very thin. | 0 | 0 | 0 | 2 | 0 | 1 |
| U676 | 05-02-88 | - | No history. | 0 | 0 | 1 | 1 | 0 | - |
| U677 | 18-03-88 | J/F | Clenched feet, but able to grip, depressed, opisthotonus, unable to fly. Euthanised. Very thin. | 0 | 2 | 2 | 0 | 0 | - |
| U735 | 02-06-88 | J/M | Neurological signs. Thin. Euthanised. | 0 | 0 | 0 | 2 | 0 | 1/WD |
| U782 | 31-08-88 | J/M | Caught be cat. Clenched feet. Leg paresis. Euthanised. Thin | 2 | 0 | 0 | 1 | 1 | 1 |
| U783 | 29-08-88 | A/M | Clenched feet. Euthanised. | 0 | 0 | 2/N1 | 2/N1 | 0 | - |
| U784 | 02-09-88 | J/F | Clenched feet. Died. Very thin. | 1 | 1 | 0 | 2 | 0 | - |
| U785 | 19-09-88 | A/F | Clenched foot. Unable to fly. Eating well. Euthanised. Good condition. | 0 | 0 | 0 | 3 | 0 | 2 |
| U793 | 27-09-88 | A/M | Neurological signs. Euthanised. Thin. | 0 | 0 | 2 | 2 | 0 | - |

| | | | | | | | | | |
|---|---|---|---|---|---|---|---|---|---|
| U829 | 13-12-88 | J/M | Clenched feet. Euthanised. Good condition. | 0 | 0 | 2/N1 | 3 | 0 | - |
| U831 | 16-12-88 | A/F | Clenched feet. Euthanised. Thin | 0 | 0 | 0 | 2/N1 | 0 | WD |
| U914 | 17-03-89 | A/F | Paresis, unable to use wings. Euthanised. Thin. | 0 | 0 | 0 | 2 | 0 | - |
| U943 | 21-04-89 | A/M | Clenched feet, drooping wing. Euthanised. Good condition. | 1 | 0 | 0 | 3 | 0 | 1 |
| U1049 | 12-09-89 | J/M | Clenched feet. Very thin. | 0 | 0 | 2 | 2 | 0 | - |
| U1097 | 1989 | - | Clenched feet. | 0 | 0 | 1 | 1 | 0 | - |
| U1107 | 17-01-90 | J/F | Clenched feet. Good condition. | 0 | 0 | 1 | 2 | 2 | 2 |
| U1123 | 31-11-90 | - | Clenched feet. | - | - | - | 2 | 0 | - |
| U1418 | 23-11-91 | A/F | Paralysis. Euthanasia. Very thin. | 0 | 0 | 0 | 2 | 0 | - |
| U1391 | 18-12-90 | A/F | Clenched feet. Euthanised. Good condition. | 0 | 0 | 0 | 3 | 3 | 0/WD |
| U1902 | 19-01-93 | A/M | Clenched foot. Tremor. Euthanised. Thin. | 0 | 0 | 1/N1 | 1 | 2 | 1/WD |
| U1903 | 19-11-93 | A/M | Clenched foot. Euthanasia. Good condition. | 0 | 0 | 0 | 1/N1 | 0 | 1/WD |
| U2361 | 01-10-94 | A/F | Dog attacked. Head tilt. Lethargic. Euthanised. Thin. | 0 | 3/N2 | 3/N2 | - | - | - |
| U2408[Q] | 10-1994 | J/M | Clenched feet. Euthanised. | 1 | 1 | 1 | 0 | - | - |
| U2409[Q] | 10-1994 | - | Clenched feet. Euthanised. | 0 | 2 | 2 | - | - | - |
| U2463 | 1995 | - | Clenched feet. Weak. Walking on hocks. | 0 | 1 | 0 | 3 | 0 | WD |
| U2464 | 1995 | A/- | Clenched feet. Lateral recumbency. | 0 | 0/WD | 0 | 2 | 0 | WD |
| U2842[Q] | 18-10-96 | - | Clenched feet. Euthanised. | 0 | 3 | 3 | 3 | 0 | WD |
| U2945 | 08-10-97 | A/M | Clenched feet. Unable to fly. Euthanised. Good condition. | 0 | 0 | 0 | 2 | 0 | - |
| U2948[Q] | 1997 | - | Clenched feet. Euthanasia. | 0 | 1 | 1 | 2 | 0 | WD |
| U3099 | 30-09-98 | J/M | Head tilt, ataxia. Euthanised. Very thin. | 2 | 2 | 2 | 1 | 0 | - |
| 2989.1 | 26-07-02 | A/F | Clenched feet, head tremor, ataxic. Euthanised. Thin. | 0 | 1 | 0 | - | - | - |
| 3153.1 | 23-10-02 | A/F | Paresis, marked intention tremor. Euthanised. Good condition. | 0 | 1 | 1 | 1/WD | 0 | - |
| 4575.1 | 01-03-05 | A/F | Clenched foot, unable to fly, dyspnoea. Euthanised. Good condition. | 0 | 1 | 0 | - | - | - |
| 5604.1 | 23-01-07 | A/M | Clenched foot, balancing on hocks and wings. Euthanised. Thin. | 2 | 0 | 0 | - | - | - |
| 5789.1 | 21-05-07 | A/F | Depressed, immobile. Euthanised. Emaciated. | 0 | 2 | 2 | - | - | - |
| **Total tissues with non-suppurative inflammation/total number of tissues examined** | | | | **54** | **54** | **54** | **49** | **47** | **19** |
| **Total number classified in each grade of severity- 0/1/2/3** | | | | **42/7/5/0** | **28/10/12/4** | **24/9/17/4** | **7/12/23/7** | **41/1/3/1** | |

All cases originate from NSW except Queensland cases, which are denoted by [Q].
A - Adult, J - Juvenile, F- Female, M- Male, – - No data.
Non-suppurative inflammation graded on a scale of 0-4 in ascending severity. Necrosis (N) graded on a scale of 0-4 in ascending severity. Wallerian Degeneration (WD) indicated when present.

**Supplementary Table 2** Presentation and pathology of passerines with myocardial degeneration and myocarditis.

| Species | Case ID | Date | Location | Signalment | Presentation | Myodegeneration/ inflammation | Cardiac myodegeneration/ inflammation | CNS inflammation | Hepatic necrosis/ inflammation | Enteric necrosis/ inflammation | Pancreatic inflammation | Vasculopathy |
|---|---|---|---|---|---|---|---|---|---|---|---|---|
| Magpie | 3630.3 | 1/08/2003 | Budgewoi | J/M | Found dead. Good condition. Mass mortality (n=35). | 0/0 | 0/1 | 0 | 0/0 | 0/2 | 0 | 1 |
| Currawong | 3630.4 | 1/08/2003 | Budgewoi | A/M | Found dead. Good condition. Mass mortality. | 0/0 | 0/1 | 0 | 0/0 | 0/0 | 0 | 0 |
| Currawong | 3667.1 | 27/08/2003 | Balmoral | A/F | Weak, paresis. Euthanised. Good condition. Mass mortality (n=4). | 3/3 | 1/1 | 1 | 0/2 | 0/0 | 0 | 0 |
| Magpie | 3687.1 | 1/08/2003 | Canley Heights | A/M | Found dead. Diarrhoea. | 0/0 | 0/1 | 0 | 0/0 | 0/0 | 0 | 0 |
| Magpie | 5103.1 | 20/02/2006 | Tuggerah | J/M | Paresis, weak, alert, diarrhoea. Euthanised. Thin. Mass mortality (n=23). | 0/1 | 0/1 | 1 | 0/0 | 0/1 | 0 | 1 |
| Magpie | 5103.2 | 15/02/2006 | Tuggerah | J/M | Paresis, dyspnea. Euthanised. Thin. Mass mortality. | 0/0 | 2/0 | 1 | 0/0 | 0/2 | 0 | 1 |
| Magpie | 5103.3 | 16/02/2006 | Kulnura | A/F | Recumbent, weak, flaps wings, alert. Euthanised. Good condition. Mass mortality. | 0/1 | 0/1 | 1 | 0/0 | 0/2 | 0 | 2 |
| Magpie | 5103.4 | 23/02/2006 | Wyong | A/F | Weak, recumbent, alert. Good condition. Mass mortality. | 1/2 | 0/2 | 2 | 0/0 | 0/0 | 0 | 1 |
| Magpie | 5103.5 | 12/02/2006 | Kincumber | A/M | Recumbent, weakness, flaps wings, alert. Euthanised. Thin. Mass mortality. | 1/2 | 1/1 | 1 | 0/0 | 0/0 | 0 | 2 |
| Magpie | 5103.6 | 16/02/2006 | Woy Woy | A/F | Weak, aggressive and alert. Euthanised. Thin. Mass mortality. | 0/1 | 2/1 | 0 | 0/1 | 0/0 | 0 | 1 |
| Magpie | 5103.7 | 16/02/2006 | Umina | A/M | Weak, aggressive and alert. Euthanised. Thin. Mass mortality. | 0/2 | 1/1 | 0 | 0/0 | 0/0 | 0 | 1 |
| Magpie | 5103.8 | 14/02/2006 | Umina | A/M | Weak. aggressive and alert. Euthanised. Thin. Mass mortality. | 0/0 | 1/1 | 1 | 0/0 | 0/0 | 0 | 1 |
| Magpie | 5103.9 | 25/02/2006 | Bateau Bay | A/F | Died in Transit. Euthanised. Good condition. Mass mortality. | 0/1 | 2/0 | 0 | 0/1 | 0/0 | 2 | 2 |
| Magpie | 5103.10 | 25/02/2006 | Bateau Bay | A/M | Paresis, moribund. Euthanised. Thin. | 1/1 | 1/1 | 1 | 0/0 | 0/0 | 0 | 1 |
| Magpie | 5103.11 | 24/02/2006 | Wyoming | A/M | Weakness. Euthanised. Thin. Mass mortality. | 0/1 | 1/1 | 1 | 0/1 | 0/1 | 0 | 0 |
| Raven | 5103.12 | 8/03/2006 | Kulnura | J/M | Recumbent, clenched feat, weak, slow righting reflex. Thin. Mass mortality. | 0/0 | 1/0 | 2 | 0/0 | 2/2 | 0 | 0 |
| Magpie | 5104.1 | 21/02/2006 | Fairlight | J/M | Weakness. Parasites. Euthanised. Very thin. | 0/1 | 0/0 | 1 | 0/0 | 0/1 | 0 | 0 |
| Currawong | 5109.1 | 22/02/2006 | Kingsford | J/M | Weak, flaps wings well, alert. Euthanised. | 0/0 | 0/1 | 0 | 0/0 | 0/0 | 0 | 1 |
| Currawong | 5119.1 | 26/02/2006 | Coogee | A/M | Weak, alert. Died. Very thin. | 2/1 | 1/0 | 0 | 0/2 | 0/0 | 0 | 0 |
| Magpie | 5120.1 | 27/02/2006 | Matraville | A/M | Weak, alert. Died. Thin. | 0/1 | 1/1 | 1 | 0/0 | 0/0 | 0 | 1 |
| Magpie Lark | 5127.1 | 8/03/2006 | Leichardt | A/F | Weak. Euthanised. Thin. | 0/0 | 2/0 | 0 | 0/1 | 0/0 | 0 | 1 |
| Currawong | 5134.2 | 12/03/2006 | Kirrawee | A/M | Weak, slow righting, aggressive and alert. Euthanised. Thin. | 2/2 | 0/1 | 1 | 0/0 | 0/0 | 0 | 0 |
| Raven | 5135.1 | 14/03/2006 | Killarney Height | A/M | Paresis, aggressive, alert. Thin. | 1/0 | 2/0 | 1 | 0/0 | 1/1 | 0 | 0 |
| Raven | 5168.1 | 31/03/2006 | Balmoral | A/M | Recumbent, weak, dyspnoeic, bloody diarrhoea. Thin. | 0/0 | 0/1 | 1 | 2/1 | 0/1 | 0 | 1 |
| Currawong | 5519.1 | 25/11/2006 | Kensingtion | J/- | Weak, ataxia, slow righting. Died. Good condition. | 1/1 | 1/1 | 0 | 0/1 | 1/0 | 0/1 | 1 |
| Currawong | 5519.2 | 25/11/2006 | Kensingtion | J/- | Paresis, weakness, poor neck control. Euthanised. Good condition. | - | 0/1 | 0 | 0/1 | 0/0 | 0 | 1 |
| Currawong | 5618.1 | 24/11/2006 | Cromer | A/- | No history. Died. Good condition. | - | 0/2 | 0 | 2/2 | 0/0 | 0 | 0 |
| Currawong | 5618.2 | 23/11/2006 | Cromer | A/- | No history. Died. Good condition. | 0/2 | 1/2 | 0 | 2/2 | 0/0 | N/A | 0 |
| Currawong | 5606.1 | 23/01/2007 | Ashfield | A/M | Found dead. Good condition. Mass. | - | 1/1 | 0 | 1/1 | 0/0 | 0 | 0 |

| | | | | | | | | | | | | |
|---|---|---|---|---|---|---|---|---|---|---|---|---|
| Magpie | 5624.1 | 2/02/2007 | Randwick | A/M | Paralysis except head, alert. Died. Good condition. | 2/0 | 0/1 | 0 | 1/1 | 0/0 | 0 | 0 |
| Currawong | 6736.1 | 2/02/2009 | Manly | A/F | Paresis, flaps wings and eats. Euthanised. Thin. | - | 0/1 | 1 | 0 | 0/0 | 1 | 0 |
| Currawong | 6751.2 | 12/02/2009 | Oyster Bay | A/M | Paresis. Weak. Gurgling. Died. Thin. | 1/2 | 2/2 | 1 | 0/1 | 0/0 | 1 | 0 |
| Magpie | 6739.1 | 5/01/2009 | Hurstville | J/M | Weak, hock siting, withdrawal reflexes, diarrhoea. Euthanised. Thin. | 0/1 | 0/1 | 2 | 0/0 | 0/0 | 0 | 1 |
| Raven | 7139.1 | 21/08/2009 | Fairlight | J/- | Weak, ataxic, head tilt, circling. Euthanised. Thin. | 0/0 | 1/0 | 3 | 2/0 | 0/0 | 0 | 0 |
| Raven | 7200.1 | 1/10/2009 | Balgowlah | J/- | Ataxic, odd head movements, unable to perch, but eating and alert. Euthanised. Thin. | 0/0 | 0/1 | 2 | 2/0 | 0/2 | 1 | 2 |
| Currawong | 7886.1 | 22/11/2010 | Mosman | J/M | Nestling - fell out of nest. Euthanised. Good condition. | 0/0 | 3/3 | 0 | 2/2 | 0/0 | 1 | 2 |
| Currawong | 7886.2 | 22/11/2010 | Mosman | J/- | Nestling - fell out of nest. Euthanised. Thin. | 0/0 | 3/3 | 0 | 2/2 | 1/2 | 2/2 | 2 |
| Magpie | 7912.1 | 1/12/2010 | Balmoral | J/- | Nestling - fell out of nest, parasites. Euthanised. Emaciated. | 0/0 | 0/1 | 0 | 0/1 | 0/2 | 0 | 2 |
| Magpie | 8536.1a | 18/02/2012 | Aitkenvale QLD | A/F | Sudden death in rehabilitation care. Good condition. | 0/0 | 0/1 | 1 | N/A | 0/0 | 0 | 0 |
| Magpie | 8536.1b | 18/02/2012 | Aitkenvale QLD | A/F | Sudden death in rehabilitation care. Good condition. | 0/0 | 0/2 | - | 0/2 | 0/0 | 0 | 0 |
| Figbird | 8599.2 | 24/04/2012 | Haberfeld | A/F | Found dead. Not toxins. Good condition. Mass mortality (n=13). | 0/0 | 0/2 | 0 | 0/0 | 0/2 | 0 | 1 |
| Magpie | 9585.2 | 29/10/2013 | Nowra | J/M | Fledglings in care, anorexia, lethargy, parasites. Died. Thin. Mass mortality (n=21). | 1/2 | 2/2 | 1 | 1/1 | 2/2 | 2 | 2 |
| Magpie | 9585.3 | 29/10/2013 | Nowra | J/F | Fledglings in care, anorexia, lethargy, parasites. Died. Thin. Mass mortality. | 0/2 | 0/2 | 1 | 0/1 | 0/2 | 0 | 2 |
| Currawong | 9586.1 | 29/10/2013 | Mosman | J/M | Nestling - fell out of nest. Euthanasia. Good condition. | 0/2 | 3/2 | 2 | 2/0 | 0/0 | 1 | 1 |
| Magpie | 9900.1 | 10/05/2014 | Avalon | A/F | Weakness. Not toxins. Good Condition. Mass mortality (n=18). | 0/1 | 0/3 | 0 | 0/0 | 1/2 | 2 | 2 |
| Magpie | 9900.3 | 10/05/2014 | Avalon | J/M | Weakness. Not toxins. Good Condition. Mass mortality. | 0/1 | 0/1 | 0 | 0/0 | 0/2 | 0 | 0 |
| Raven | 10444.1 | 19/02/2015 | Mosman | J/F | Weak, recumbent, head curled under body, dyspneoa. Thin. | 0/0 | 1/0 | 2 | 0/0 | 0/2 | 0 | 0 |
| Magpie | 10592.1 | 20/05/2015 | Oak Flats | A/F | Weak. Mass mortality. Euthanasia. Fenthion detected 3/4 liver samples. Good condition. Mass mortality (n=17). | 0/0 | 0/2 | 0 | 2/0 | 2/2 | 0 | 2 |
| Magpie | 10592.2 | 20/05/2015 | Oak Flats | A/M | Weak. Mass mortality. Euthanasia. Fenthion detected 3/4 liver samples. Good condition. Mass mortality (n=17). | 0/0 | 0/2 | 0 | 2/2 | 0/2 | 1 | 2 |

A – Adult, J – Juvenile, F- Female, M- Male, - signifies not available for examination.

Myodegeneration, inflammation, necrosis, and vasculopathy graded on a scale of 0-4 in ascending severity.

Locations are within NSW unless otherwise specified. Shaded cases represent clusters or epizootics.

**Supplementary Table 4**. Primers used to recover the full length genomes of the viruses identified here.

| **Primer IDs** | **Forward primer sequence (5'-3')** | **Reverse primer sequence (3'-5')** | **Target region** | **Amplicon size (bp)** |
|---|---|---|---|---|
| Polyoma S1 | GGTAGTGACGGTATTTTGAGGTC | CAAGTAGATTCACCTCAGCTCTC | 220-1815 | 1605 |
| Polyoma S2 | GCTGCTATAACTGCTTTAGAAGGT | CAGAAAGATAAAGCCCGTCACC | 1050-2587 | 1538 |
| Polyoma S3 | GACCCTACTCTTAAAGCCAGACT | TTCTCACGGGTGCCAGTTTAA | 2368-3571 | 1204 |
| Polyoma S4 | ATGTGGAATCTCGGTGAAAGTC | TGAAACGCCTTAATTGCCTGA | 3223-4968 | 1736 |
| Polyoma S5 | AGTGTGGATCTTGCAGCTTCA | GTCACTATATGCCTGTAAATCGTCC | 4896-815 | 1024 |
| Paramyxo S1 | CACAGACTATGATAAGTTGCAGG | GTTGGTTACTGCTTGGATCATG | 137-2125 | 1989 |
| Paramyxo S2 | GCAAATCCCAATTCCAGCAAC | CTTCACATGAGGATTGATGGAGG | 1396-3343 | 1948 |
| Paramyxo S3 | TGAATGCTGGTCTCAATGAATGG | CTCTTGGAGAAGGGTTTGTTGAC | 3272-5308 | 2037 |
| Paramyxo S4 | GAACAGAAGGAAAGAGTGATTACG | AGAGGTCTGGATATTATGGACGA | 4953-6653 | 1701 |
| Paramyxo S5 | GAATCATGGAGGAGGTTGACC | GCCCACAATGAGCCTCTAAT | 6572-8725 | 2154 |
| Paramyxo S6 | GCACTGTCTAATTCCCCTGATT | GATCTGTGTACATAAGGACCATCT | 7988-10547 | 2560 |
| Paramyxo S7 | CGTCTTAAATTCCATTACTGCGC | CAAGTCATGTGGTGGGTTGTAT | 9973-12029 | 2057 |
| Paramyxo S8 | GCAACTCATATCTGTGACTTCTTC | GGATCTCAAATGGCAAGTCATG | 11546-13613 | 2068 |
| Paramyxo S9 | TCATGGCGTCCGTTATTACAAG | AAGTCTTGGATGAAACCCTTGG | 13332-15487 | 2156 |
| Paramyxo S10 | GCACCTATTCATGAGTTGTTGAC | GTTTCACTTCATGTCTAGTCAGAG | 14847-16512 | 1666 |

| | | | | |
|---|---|---|---|---|
| Adv_hexon | CACATAGCGGGTCTTTAGCAAC | CAGTTGCAAAAGGAGTTCTGAAG | 14550-16376 | 1827 |
| Adv_pol | TGTCTTCTTTGGTAGCGAAAC | ATTCTTCTTGCGAGCCAGAT | 4557-5864 | 1308 |

**Supplementary Table 3**. PCR primers used to amplify viral sequences from bird tissues.

| **Primer IDs** | **Forward primer sequence (5'-3')** | **Reverse primer sequence (3'-5')** | **Target** | **Amplicon size (bp)** |
|---|---|---|---|---|
| Clenched claw syndrome | | | | |
| BRDV-cap-F/R | CTTGTAGTGGDATCCABCCG | GTGGAGCACCTCTVACTGC | Circovirus-cap | 596 |
| MET011/12 | AACCAGATCCTCGGTATACC | TGGACCTCTCTTGTGATAGC | Paramyxovirus | 301 |
| MET013/14 | ATAATGACATGCGATGTGCTC | CTAATGCTAAGATCAAGTCCTACC | Hepatovirus | 220 |
| MET150/151 | TGCTTAATCCAATTTTTAATCCAGA | CATCCTGGACACATTGTTATTCA | Adenovirus-pol1 | 113 |
| MET154/155 | TGATTTTATTCTGTGATTTCCATG | GGTGACACTGACAGCTTATT | Adenovirus-pol2 | 78 |
| MET181/182 | CTGATAATGGAGCTTATACAACGG | TGTGTATCCAATGCAGTATACGG | Parvovirus | 248 |
| Black and white bird diseases | | | | |
| MET029/030 | CCTTTCCTGATCAGTGTTACC | TATTGAGATGGACTGGACTCG | Astrovirus | 195 |
| MET009/010 | GTTGACGATCACATCAAAGTG | ATTCAGACAGATAGGTATCAACC | Picornavirus | 220 |
| MET031/032 | TAGATCTGCAGGGTCTAGTG | CCATCCTGATCAAGTCTGG | Polyomavirus | 147 |
